# Information Theory of Composite Sequence Motifs: Mutational and Biophysical Determinants of Complex Molecular Recognition

**DOI:** 10.1101/2024.11.11.623117

**Authors:** Elia Mascolo, Ivan Erill

**Affiliations:** Departament d’Enginyeria de la Informació i de les Comunicacions, Universitat Autònoma de Barcelona, Bellaterra, Spain; Department of Biological Sciences, University of Maryland Baltimore County, Baltimore, MD, USA

**Keywords:** biological information theory, sequence motifs, molecular machines, evolution, mutational robustness, transcriptional regulation, energy dissipation

## Abstract

The recognition of nucleotide sequence patterns is a fundamental biological process that controls the start sites of replication, transcription and translation, as well as transcriptional and translational regulation. Foundational work on the evolution of biological information showed that the amount of information encoded in the target nucleotide sequence patterns, a quantity named *R*_*sequence*_, evolves by natural selection to match a predictable quantity called *R*_*frequency*_. In this work, we propose a generalization of this canonical framework that can describe composite sequence motifs: motifs composed of a series of sequence patterns at some variable (not necessarily conserved) distance from each other. We find that some information can be encoded through the conservation of the distance between sequence patterns, a quantity we named *R*_*spacer*_, and that - to be functional - biological systems require the sum of *R*_*sequence*_ and *R*_*spacer*_ to be constant. We empirically validate our mathematical results through evolutionary simulations. We apply this general framework to demonstrate that the pre-recruitment of regulatory complexes to target sites has intrinsic advantages over in situ recruitment in terms of energy dissipation and search efficiency, and that realistic values of protein flexibility co-evolve with the target composite motifs to match their spacer size variability. Lastly, we show that the relative advantage of encoding information in sequence patterns or in spacers depends on the balance between nucleotide substitutions and insertions/deletions, with known estimates for the rates of these mutation types favoring the evolution of composite motifs with highly conserved spacer length.

## 1. Introduction

Many cellular processes depend on the binding of protein or RNA complexes to specific locations within the genome. These macromolecules typically recognize short (5-50 bp) contiguous sequence patterns, or motifs. Given a set of aligned binding sites targeted by one of these macromolecular recognizers, one can compute the relative frequency of each base at each position of the motif, which is typically represented as a position-specific frequency matrix (PSFM) (Fig. 1). Assuming that each position contributes independently to binding, a fundamental result in molecular information theory was the derivation of *R*_*sequence*_ as a metric for the amount of information encoded in the motifs targeted by macromolecular recognizers [1].

**Figure 1:**
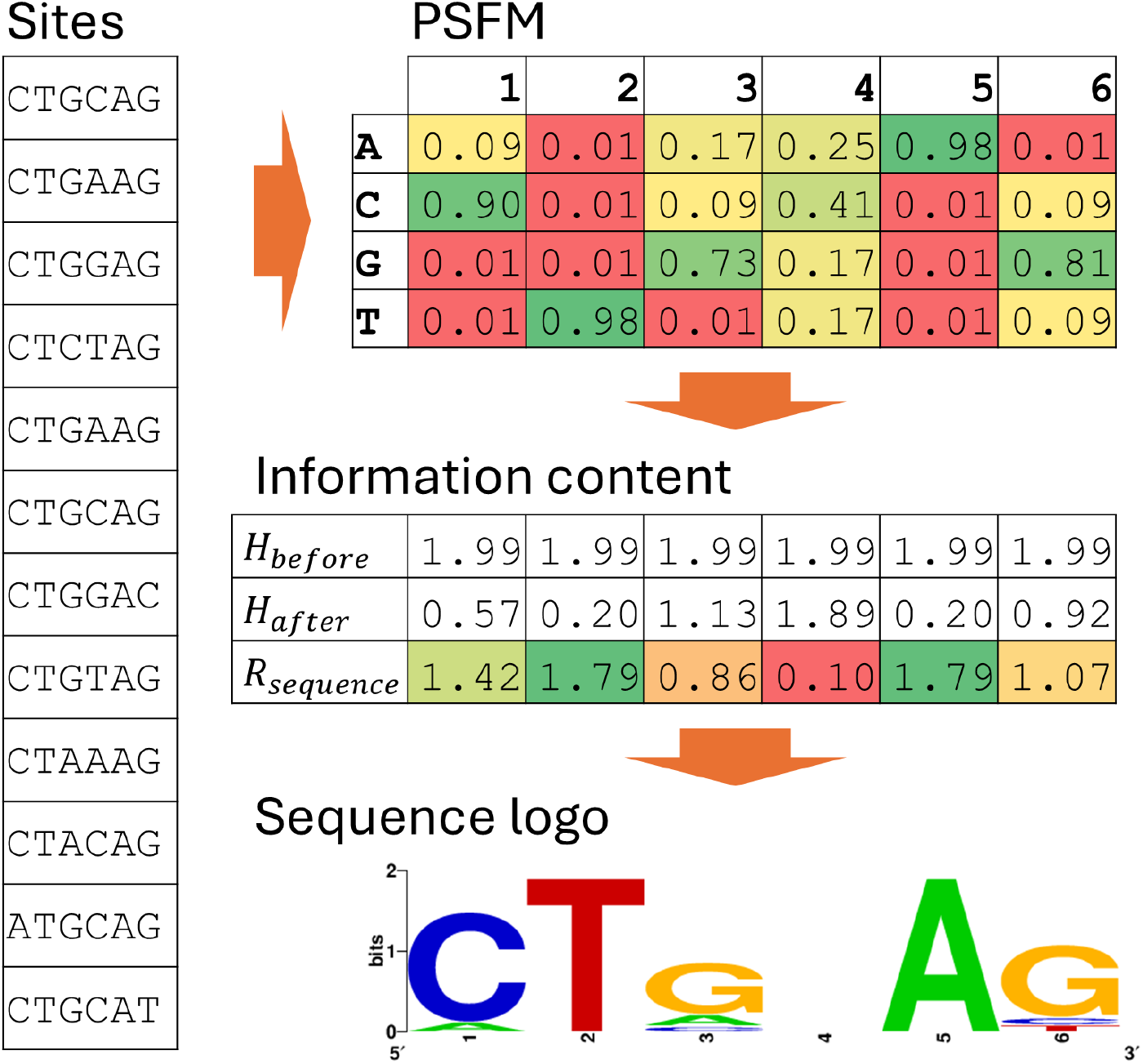
Schematic derivation of the position-specific frequency matrix (PSFM) and the information content (*R*_*sequence*_) for a set of sites bound by a fictional recognizer operating on the *Escherichia coli* genome. Frequency estimation uses additive smoothing with smoothing parameter α = 0.1 [2]. A priori entropy values are computed using the *E. coli* genome sequence (NC 000913.3) base frequencies (A/T: 24.6%; C/G: 25.4%). The sequence logo was generated using the WebLogo service [3].

In the absence of additional information, our uncertainty over the bases occupying any position of a fragment of length L of a genomic sequence can be measured using Shannon’s entropy [4] as

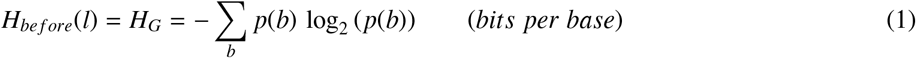

where *p*(*b*) are the genomic frequencies of each DNA base *b* ∈ {*A, C, G, T*}. Once we know that a recognizer is specifically bound to a target sequence fragment, however, our uncertainty over the bases occupying each position decreases and is dictated by the specificity of the binding motif at each position:

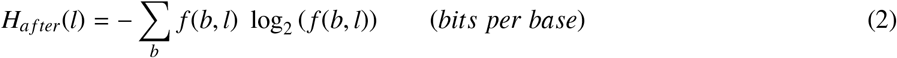

where *f* (*b, l*) is the frequency of DNA base *b* at position *l* of the motif. It follows that our observation of the recognizer binding to the sequence fragment can be formally treated as an information gain. Using Shannon’s definition [4], the amount of information gained on the bases occupying each position of the fragment can be expressed as the difference between a priori and a posteriori entropies:

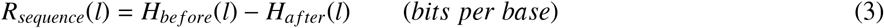

Assuming positional independence, the information content of the entire recognizer-binding motif is classically expressed as:

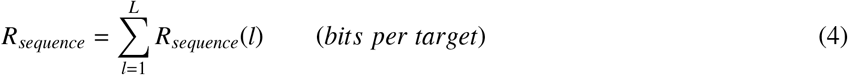

Another fundamental result in molecular information theory was the derivation of *R*_*frequency*_, which measures the amount of information required by a recognizer to fulfill its function [1]. The task of the recognizer is to identify *γ* binding sites out of *G* genomic positions. Therefore, the target recognition process is a reduction in positional uncertainty. Our initial uncertainty over the position of a recognizer targeting *γ* sites in a circular genome of length *G* is the a priori positional entropy:

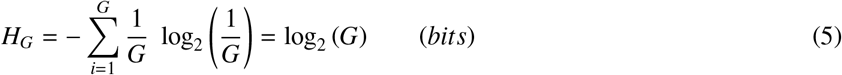

Once the recognizer binds specifically to one of its *γ* target sites, however, our uncertainty is reduced to the a posteriori positional entropy:

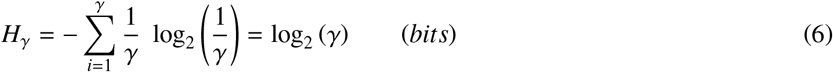

The information we gain on the location of the recognizer can again be expressed as the difference between a priori and a posteriori entropies:

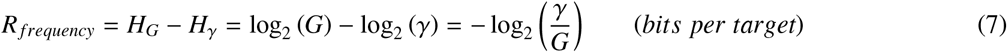

which is simply a function of *γ*/*G*, the frequency of target sites in the genome.

The *R*_*sequence*_ metric defines the information encoded by the target binding sites through nucleotide conservation, whereas *R*_*frequency*_ determines the information that is required for a recognizer to specifically bind one of its *γ* genomic targets. Hence, it follows that for the recognizer to be operational in its genomic environment, *R*_*sequence*_ must be greater or equal to *R*_*frequency*_. Since the encoding of *R*_*sequence*_ in target sites is driven and maintained by natural selection, only the information required by the recognizer to perform its function will be preserved, and it is therefore expected that *R*_*sequence*_ be approximately equal to *R*_*frequency*_. This result is supported by experimental evidence on transcription factor- and ribosome-binding motifs [1] and was elegantly established by the use of evolutionary simulations, in which both recognizer-binding sites and the sequence specificity of the recognizer are allowed to co-evolve freely [5]. In these simulations, the sequences of designated recognizer-binding site locations evolve to reach *R*_*sequence*_ values close to the predicted *R*_*frequency*_ value.

The formalization of recognizer-DNA binding events as information processes enables the interrogation of these systems under the mathematical framework of information theory, and the subsequent derivation of the near equality between *R*_*frequency*_ and *R*_*sequence*_ provides important insights into the evolutionary dynamics of recognizers and their genomic targets.

In many biological systems, however, recognizers bind to DNA or RNA synergistically, jointly targeting multi-element, or *composite*, motifs. The information that prokaryotic ribosomes recognize to initiate translation in mRNA, for instance, is encoded as a composite motif defined by two conserved sequence patterns, the Shine-Dalgarno sequence and the initiation region, separated by a variable spacer [6]. Similarly, the complex of nucleotide-binding proteins and small nuclear RNAs forming the eukaryotic spliceosome identifies exon-intron borders in precursor mRNA by recognizing a composite motif composed of three sequence patterns (donor, branchpoint and acceptor sites) located at variable distances from each other [7]. Composite motifs are also a common feature in transcriptional regulatory systems. Transcription initiation in eukaryotes is typically regulated through the combinatorial binding and interaction of multiple transcription factors in regulatory modules [8], and in a substantial number of these interactions, the order and relative distance between binding sites plays an important role in regulatory activity [9, 10, 11]. Well-known instances of this phenomenon are the composite binding sites of heterodimers formed by different nuclear receptors with retinoid X receptors [12] or the organization of the viral interferon-β enhanceosome [13]. In bacteria, composite motifs have been reported for both homologous and heterologous multimers, such as transcription factors of the PrrA, IclR and MocR families [14, 15, 16] or the CytR-CRP complex [17]. In all these cases, composite motifs combine multiple sequence patterns located at variable distances from each other and cannot therefore be adequately modeled as contiguous sequence patterns, hindering their formal analysis using information theory.

In this work, we generalize the information theory framework of sequence motifs to encompass composite motifs consisting of an arbitrary number of sequence patterns located at variable distances from each other. We derive a formal expression for the information encoded by a composite sequence motif, which includes contributions from each constituent sequence motif (*R*_*sequence*_) and their intervening distances (*R*_*spacer*_). We establish that the predicted amount of information required to locate composite binding sites (*R*_*frequency*_) can be arbitrarily distributed among sub-motifs and their relative spacing constraints, and we use evolutionary simulations to validate the predictions made by the theory. Furthermore, we leverage this new framework to explore the informational and thermodynamic requirements of recruitment and pre-recruitment strategies for DNA-binding protein complexes, and to address the preference for conserved spacer length in the dyad-motifs of dimeric bacterial transcription factors. Our results show that pre-recruited recognizers can dissipate less energy per target recognition event compared to recognizers that are recruited on DNA, and that biases in substitution vs. insertion/deletion mutation rates are sufficient to explain the preference for conserved spacer lengths in bacterial transcription factor binding motifs.

## 2. Results

### 2.1. An information theory framework for composite sequence motifs

In this work, we generalize the classical theory of sequence motifs to make it applicable to composite motifs, defined as a series of sequence motifs, each at a variable distance (in base pairs) from its adjacent sequence motifs. In the context of transcriptional regulatory systems, composite motifs can be used to model those signals in regulatory DNA regions that are the targets of a combination of independent TFs or of a flexible regulator with multiple DNA-binding domains.

The simplest, non-trivial case is a composite motif with two sequence patterns, each bound specifically by a sequence-recognition element, or recognizer. This would be the case, for instance, of a transcriptional regulatory system that requires the binding of two regulatory proteins at two target sites that are at some distance (in base pairs) from each other. We will use this case of a dimeric motif, hereafter referred to as a *dyad*, to provide an intuition for each result, before generalizing to the case of a generic composite motif of *n* elements.

#### 2.1.1. Redefinition of R_frequency_

Let us consider the uncertainty about the state of the dyad-based system, which is composed of the genome and the two recognizers: *A* and *B* (*n* = 2). The entropy before target recognition is *H*_*before*_ = log_2_ (*G*^2^) bits, because the system can be in approximately *G*^2^ possible states (since both recognizers can be bound to one of the *G* possible genomic positions, but overlaps may be sterically impaired). If *γ* is the number of dyad targets (the number of composite motif instances to be recognized), the entropy after target recognition is *H*_*after*_ = log_2_ (*γ*) bits. For consistency with the classic theory, we call the reduction in positional uncertainty following target recognition *R*_*frequency*_, and we obtain:

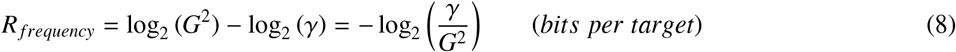

In general (*n* ≥ 1), the number of states is approximately *G*^*n*^, since each recognizer can be bound to one of the *G* possible genomic positions, and

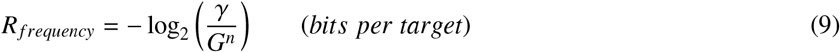

where *n* is the number of elements in the composite motif. The classical definition of *R*_*frequency*_ (Eq. 7) can now be seen as the special case when *n* = 1 (the motif is a single sequence pattern).

#### 2.1.2. Definition of R_spacer_

Let us consider again the case of a dyad-based system (*n* = 2), with *γ* dyad targets. Each of the *γ* dyad targets contains one sequence motif targeted by recognizer *A* and one targeted by recognizer *B*. Thus, *γ* is also the number of target sites for each recognizer element. Within each dyad target, the distance in base pairs between the *A* site and the *B* site (i.e., the size of the *spacer*) may be different and is distributed according to a random variable *D*. In such a system, the two recognizers may identify their correct sites within each dyad target independently (using their *R*_*sequence*_ information). Alternatively, they may exploit any expected regularity in the distance separating the *A* site and the *B* site. In other words, a regularity in *D* can provide information for target recognition. In the complete absence of such a source of information, the a priori uncertainty about the relative distance between the two recognizers would be log_2_ (*G*) bits, because (assuming genome circularity) there are *G* possible distance values that are equally probable. Therefore, we can measure the information provided by the regularity in spacer size as:

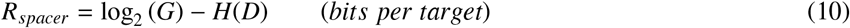

where *H*(*D*) is the entropy of *D*. The same definition applies to all the *n* − 1 spacers in a composite motif with more than two recognizers (*n* > 2).

In a dyad-based system, the spacer is maximally informative when all the dyad targets have the same spacer size, so that *H*(*D*) = 0 bits and *R*_*spacer*_ = log_2_ (*G*) bits. This special case deserves attention because a dyad with a fixed spacer could be modeled as a single sequence motif. We noted previously that the classical theory could be seen as a special case when *n* = 1. We now note that it is also equivalent to the special case when *n* > 1 but all the *n* − 1 spacers are non-variable (maximally informative). Indeed, a non-variable spacer provides log_2_ (*G*) bits of information. Therefore, for a dyad with a fixed spacer, the remaining information that must be encoded as *R*_*sequence*_ is *R*_*frequency*_ − *R*_*spacer*_ = − log_2_ (*G*2) − log_2_ (*G*) = − log_2_ (*G*) bits, which is consistent with the *R*_*sequence*_ ≈ − log_2_ (*G*) equivalence stated by the classical theory [1]. In general, in a system with *n* recognizers where all the *n* − 1 spacers are fixed, the expected amount of information to be encoded as *R*_*sequence*_ is always the amount predicted by the classical theory because

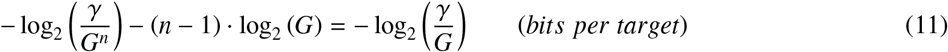

Thus, when there are no spacers (*n* = 1) or when all the spacers are maximally conserved, our theory for composite sequence motifs recapitulates the result from the classical theory. However, Eq. 11 also implies that when some spacers are not fully conserved, the total information that must be encoded as sequence patterns, 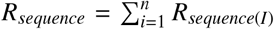, has to be higher. Therefore,

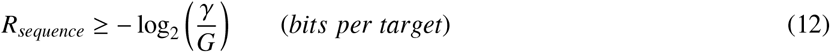

#### 2.1.3. Informational bounds

Each position in a sequence motif can encode up to 2 bits [1]. Therefore, for each sequence motif we obtain:

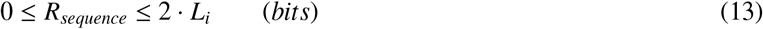

where *L*_*i*_ is the length (in base pairs) of the *i*-th sequence pattern in the composite motif and *R*_*sequence*(*i*)_ is the information encoded as sequence conservation in the *i*-th sequence pattern.

For each spacer in a composite motif, the entropy of *D* is maximal when all the *γ* targets have a different spacer size, and minimal (zero) when they have all the same spacer size (0 ≤ *H*(*D*) ≤ log_2_ (*γ*)). From the definition of *R*_*spacer*_ (Eq. 10) and substituting *H*(*D*) for the bounds above, we obtain:

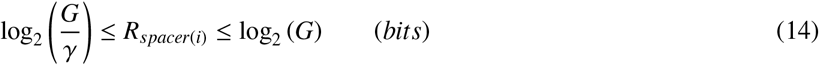

where *R*_*spacer*(*i*)_ is the information encoded as the conservation of the *i*-th spacer.

### 2.1.4. Functional bounds

Having determined what values of *R*_*sequence*_ and *R*_*spacer*_ are mathematically possible, we can now move to the problem of what values would ensure a functional biological system. Here we use the word functional to refer to a biological system such that all the targets are identified without errors. In the dyad-based system, the two recognizers (*A* and *B*) constitute a recognition complex that must recognize *γ* target dyad-sites, but none of the other possible placements of *A* and *B* on the genome. This classification task depends on the performance of each component. Each of the *γ* target dyad sites is composed of a site for recognizer *A* and one for recognizer *B*. The best that recognizer *A* can do is to correctly recognize all and only its *γ A*-sites. To achieve this task through nucleotide conservation at the *A*-sites, we need at least 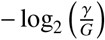 bits, as per the classical theory, and the same applies to recognizer *B*. Higher levels of conservation would not be selected because they would not provide any selective advantage [5], and would therefore be impaired by random mutations.

The fact that the *A* and *B* recognizers can both achieve perfect performances on their individual targets by means of *R*_*sequence*_ alone might suggest that a dyad-based regulator could be functional by recognizing sequence conservation only, regardless of spacer size conservation. On the contrary, even when both the sub-motifs for *A* and *B* provide 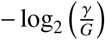 bits of information, the regulator is still missing 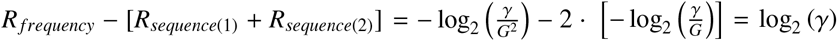 bits to properly identify the target sites of the *A-B* recognition complex. If there is only one target site (*γ* = 1), the classification task can be accomplished, since there would be log_2_ (1) = 0 missing bits. But, for *γ* > 1, additional information is required for the system to function without errors. To provide intuition for this result, it is sufficient to consider that there are *γ*^2^ dyad placements such that the *A* and *B* recognizers are both correctly placed on one of their individual sites, but only *γ* of them are the target dyad sites for the *A-B* complex (Fig. 2). Therefore, correctly distinguishing the *γ* target placements from other ways of placing recognizers *A* and *B* on their cognate sites requires log_2_ (*γ*^2^) − log_2_ (*γ*) = log_2_ (*γ*) bits, just as we found above. The only trait distinguishing those *γ*^2^ placements is the relative distance (*D*) between the two elements of the dyad. Therefore, the missing log_2_ (*γ*) bits of information must be provided by *R*_*spacer*_, via the conservation of the random variable *D*. In the previous section, we maximized the amount of information encoded in *R*_*spacer*_ and found the lower functional bound of *R*_*sequence*_ (Eq. 12). Here, by maximizing the amount of information encoded in *R*_*sequence*_, we find *R*_*spacer*_ ≥ log_2_ (*γ*) to be the lower functional bound of *R*_*spacer*_ for any individual spacer.

**Figure 2:**
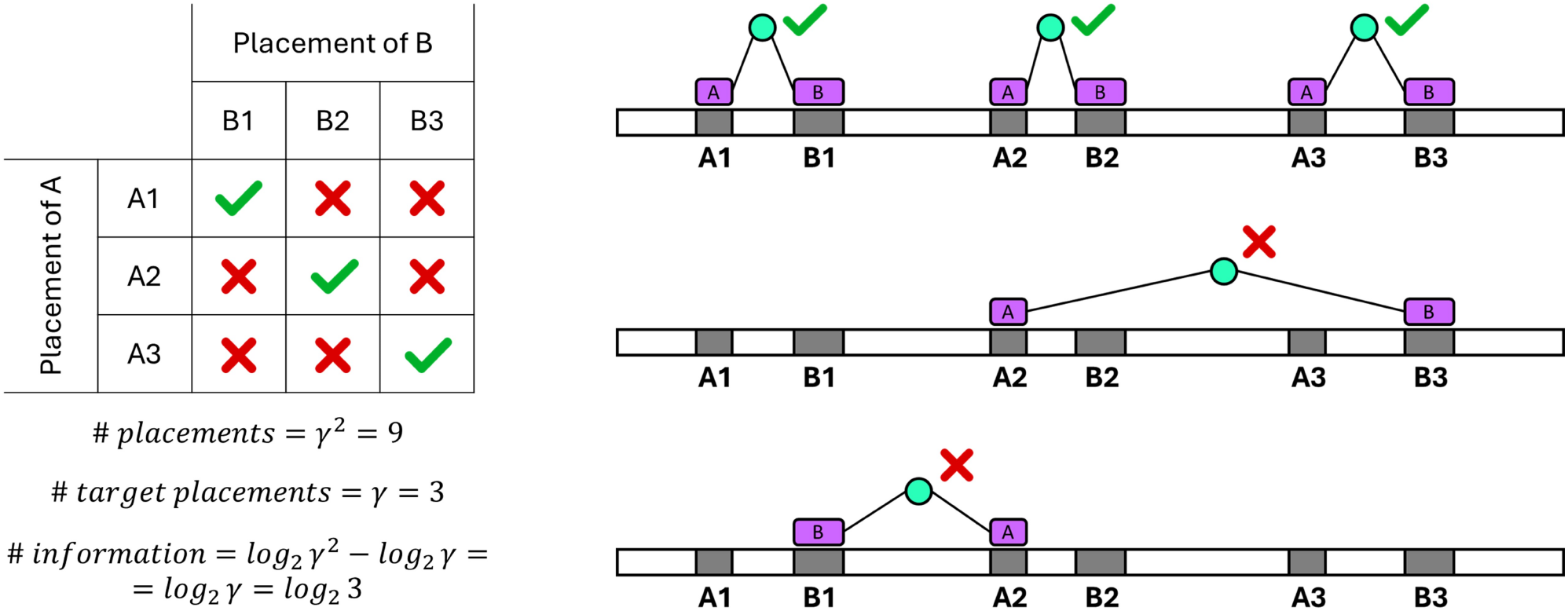
Illustration of the lower bound for *R*_*spacer*_ on a dyad-based system with two recognizers (*A* and *B*) and *γ* = 3 targets, where both recognizers can unequivocally identify their respective sites. (Left) Summary table of all the individual placements of recognizers *A* and *B*, denoting those that correspond to correct placements of the *A-B* recognizer complex on its targets. The amount of information that must be encoded in the spacer is dictated by the entropy difference between the possible *γ*^2^ combinations of placements of *A* and *B* and the *γ* correct *A-B* complex placements. (Right) Illustration of the correct recognizer complex placements on the *γ* targets, and of some of the incorrect combinations of *A* and *B* placements.

#### 2.1.5. A general theory of composite sequence motifs

By combining all the functional constraints we have derived, we obtain a general theory of composite motifs, stating that a functional system must satisfy:

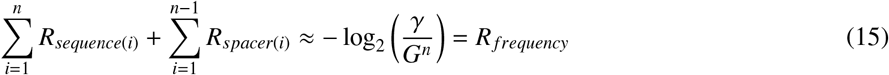

with

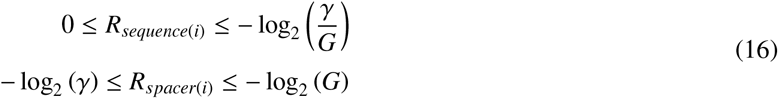

#### 2.1.6. Geometry of the solution space

Given a composite motif-based system, defined by the values of *G* (genome size), *γ* (number of targets) and *n* (number of sequence patterns in the composite motif), we can now study what encoding strategies are *functional*, in the sense that they satisfy Eq. 15. Each possible information-encoding strategy, regardless of whether or not it is functional, can be described by *n* variables of type *R*_*sequence*_ and *n* − 1 variables of type *R*_*spacer*_, which amounts to 2*n* − 1 variables in total. Thus, the possible strategies can be represented as points in a (2*n* − 1)-dimensional space, bounded by Eq. 13, 14. The *solution space* is the subset of this space that represents functional strategies. Since for a functional strategy the sum of the variables must be the constant *R*_*frequency*_ = − log_2_ (*G*^*n*^), there are only (2*n* − 1) − 1 = 2*n* − 2 degrees of freedom. Therefore, the solution space is a (2*n* − 2)-dimensional simplex. For example, when *n* = 1, Eq. 15 simplifies to the classical theory, simply stating that 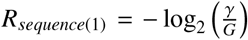. The solution is thus a single scalar because the solution space is a 0-dimensional simplex in a 1-dimensional space (i.e. a point on a line segment that represents the range of possible *R*_*sequence*_ values). When *n* = 2 the solution space becomes a 2-dimensional simplex in a 3-dimensional space (i.e. a triangle through a 3D volume), as shown in Fig. 3A.

**Figure 3:**
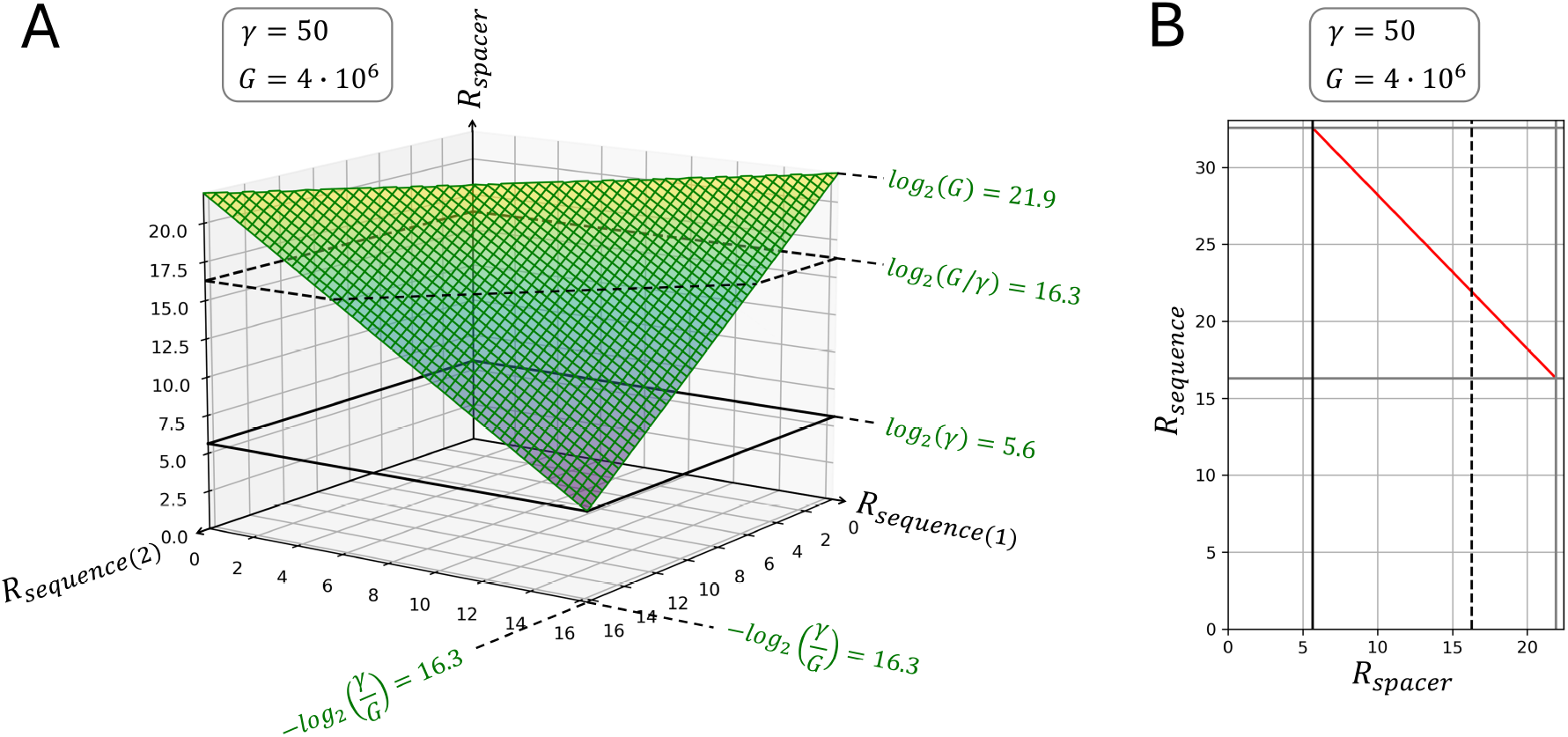
Representation of the solution space for a dyad-based system with *γ* = 50 targets in a genome of size *G* = 4 · 10^6^. The functional and informational lower bounds for *R*_*spacer*_ are represented as a solid and a dashed line, respectively. (A) Three-dimensional representation, showing the amount of information encoded in each sequence pattern (*R*_*sequence*(1)_ and *R*_*sequence*(2)_) and in the spacer (*R*_*spacer*_). The solution space is colored in green. (B) Alternative representation of the same solution space, following Eq. 17. The solution space is represented as a red line segment.

To study the relationship between *R*_*spacer*_ and *R*_*sequence*_, each information system can be characterized in a simpler way by the total information encoded via each of these two encoding strategies. In this way, our general theory can be summarized as follows:

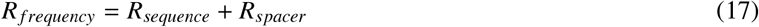

with

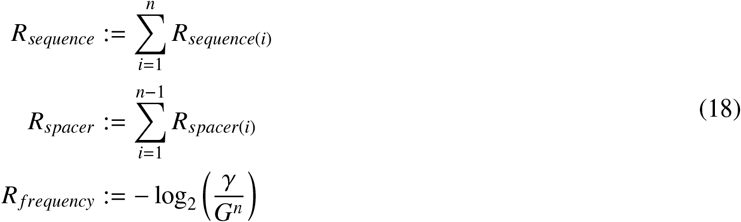

Following this description, all composite motif-based systems can be mapped onto a 2D space, regardless of the value of *n*, and their solution space is a line segment, as shown in Fig. 3B.

### 2.2. Evolutionary simulations

The classical theory of sequence motifs was empirically supported by evolutionary simulations, showing that *in silico* genomes encoding a transcription factor and containing *γ* target positions evolve sequence motif systems such that *R*_*sequence*_ ≈ *R*_*frequency*_ [5]. Here, we re-implemented a more general version of the evolutionary algorithm reported in [5] to evolve genomes encoding a recognition complex capable of identifying *γ* composite motif targets (see Methods 3.1). The genetically encoded recognition complex is composed of *n recognizers* of sequence patterns and *n* − 1 *connectors* that constrain the distance between recognizers. Each of the *G*^*n*^ possible placements is assigned a score based on how well it matches the probability distributions modeled by the recognizers (following the conventional position-specific scoring matrix (PSSM) approach; see [18, 19]) and the probability distributions modeled by the connectors (see Methods 3.2). Importantly, the theory is general and does not depend on the approach used to model recognizers and connectors. Indeed, our general theory (Eq. 17) does not assume any knowledge about the properties of the recognition complex, but only predicts a relationship between nucleotide conservation and spacer size conservation *in the targets*. The evolving “organisms” compete solely based on how good their recognition complex is at distinguishing the correct targets from the rest of the genome (they have no notion of the information theoretical metrics being tested). By setting *n* = 1, the program can be used to reproduce results from [5], showing that *R*_*sequence*_ increases until it reaches the predicted *R*_*frequency*_ value, and then it fluctuates around it (Fig. 4A). To validate the relationship between *R*_*sequence*_ and *R*_*spacer*_ described by our theory, we ran simulations with *n* = 2. We set up experiments where we controlled for spacer size conservation by specifying the spacer size in the *γ* targets, testing values of *R*_*spacer*_ that range from its minimum to its maximum (Eq. 14). In all cases, *R*_*sequence*_ evolves and converges to values such that the sum of *R*_*spacer*_ and *R*_*sequence*_ matches *R*_*frequency*_ (Fig. 4B). Thus, evolved solutions converge to the solution space that we identified for functional systems, empirically validating the predictions from our general theory (Eq. 17) (Fig. 4B and 4C).

**Figure 4:**
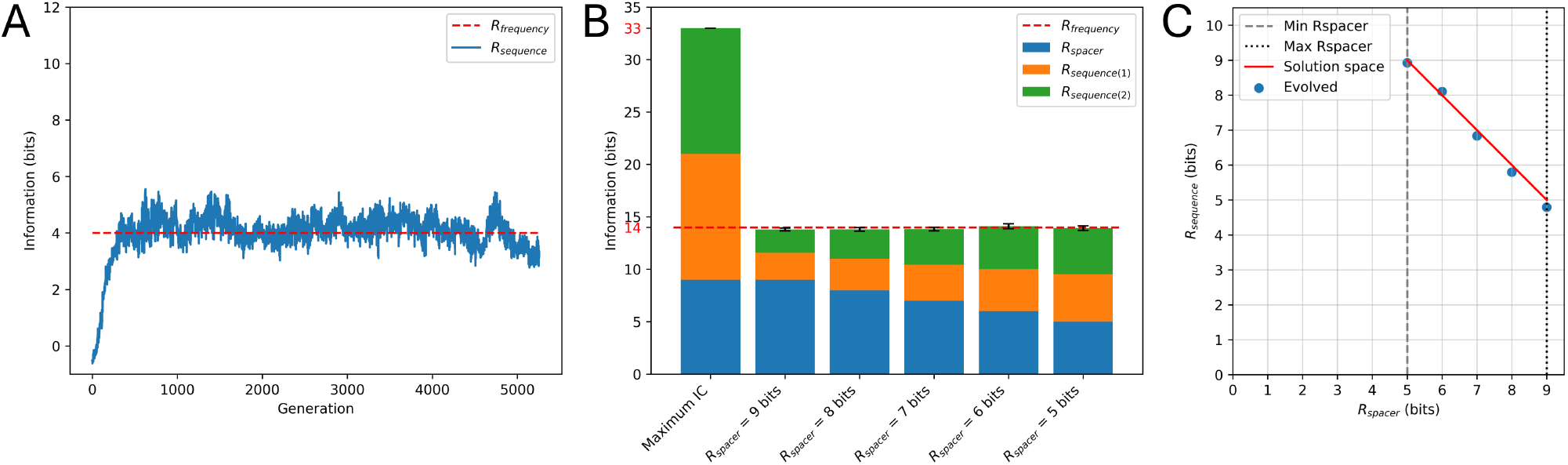
(**A**) Evolution of *R*_*sequence*_ for a single sequence motif (*n* = 1) of 6 bp, in *γ* = 16 target sites on a genome of size *G* = 256 bp. The reported *R*_*sequence*_ is the one of the organism with the highest fitness within an evolving population of 64 organisms. These were the parameters used in [5]. (**B**) Distribution of evolved information for dyad-based systems (*n* = 2) with *γ* = 16 targets on a genome of size *G* = 512 bp, with spacers encoding 9 to 5 bits of information (*R*_*spacer*_). (**C**) Mapping of evolved dyad-systems onto the theoretical solution space.

### 2.3. Energy Dissipation and Search Efficiency in Composite Motif Systems

Our theory makes an explicit distinction between *R*_*sequence*_ and *R*_*spacer*_ that can be used to study two alternative strategies for the recognition of composite motifs: the recruitment and the pre-recruitment mechanisms. In the recruitment mechanism, the *n* sequence patterns of the composite motif are recognized by *n* recognizers that search on DNA independently and that, after binding, must assemble into a recognition complex. In contrast, in the pre-recruitment mechanism, the *n* sequence patterns are recognized by a pre-assembled recognition complex, which searches for its targets as a whole (Fig. 5A). In the recruitment mechanism, the distance *δ* (expressed in bp) between a pair of recognizers has an a priori entropy *H*_*be f ore*_(Δ) = log_2_ (*G*), since each of *G* possible distances are equally likely. After complex formation, the entropy is reduced to *H*_*a f ter*_(Δ) = *H*(*D*), where *D* is the distribution of sizes in the target site spacers separating the two recognizers. In contrast, in the pre-recruitment mechanism, *H*_*be f ore*_(Δ) is dictated by the biophysical properties of the recognition complex, which will constrain the possible distances among recognizer pairs (Fig. 5B). As a result, these two mechanisms differ fundamentally in the entropy reduction that occurs during target recognition, and this impacts both the energy that is dissipated per recognition event and the efficiency of the target search process.

**Figure 5:**
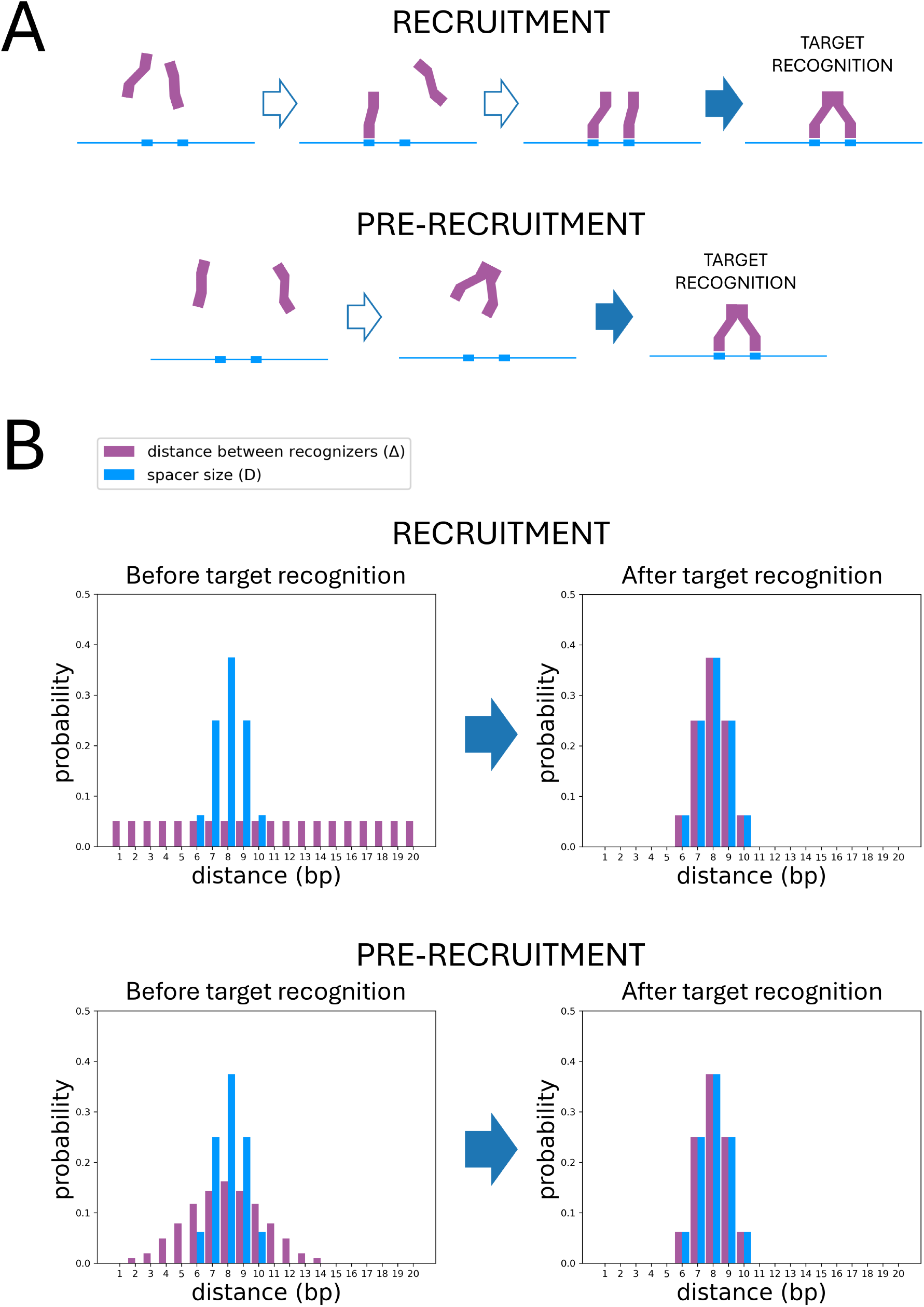
Illustration of the recruitment and pre-recruitment strategies for a dyad-based system. (**A**) Graphical representation of the recruitment and pre-recruitment strategies. (**B**) The a priori and a posteriori distribution of the distance between the two recognizers (Δ) in the recruitment and pre-recruitment strategies are shown in purple. The distribution of the spacer size (*D*) is shown in blue.

#### 2.3.1. Energy dissipation per target recognition event

A molecular entity performing target recognition on a DNA molecule can be described as a *molecular machine* [20] that discriminates *γ* targets from the genomic background. The input signals (the observed features in each placement) act like *messages* and the recognizer acts as a *receiver*. Any molecular machine gaining 1 bit of information in isothermal conditions must locally dissipate at least ℰ_*min*_ = *k*_*B*_*Tln*(2) joules per bit, where *T* is the temperature in Kelvin and *k*_*B*_ is the Boltzmann constant [21]. The precise lower bound ℰ_*min*_, also known as Landauer’s limit, has been derived independently several times in different scientific contexts [22, 23, 24, 25, 26, 27, 28, 29], and it determines an upper bound to the isothermal efficiency of molecular machines [30, 21]. Thus, proteins that recognize sequence patterns must locally dissipate at least ℰ_*min*_ *R*_*sequence*_ joules in the process of selective binding.

In the recruitment mechanism, *n* recognizers must assemble into a recognition complex after independently identifying *n* sequence patterns and collectively dissipating at least ℰ_*min*_ *R*_*sequence*_ joules. When the recognizers are not at an acceptable distance from each other, they cannot interact productively, and target recognition is not accomplished. Otherwise, they form the recognition complex, gaining information on the distance (*δ*) separating the recognizers. After target recognition, the distances between each of the recognizers are set by the size of the spacers on the sequence patterns (Δ = *D*) and, therefore, *H*_*a f ter*_(Δ) = *H*(*D*). Since *H*_*be f ore*_(Δ) = log_2_ (*G*), the reduction in entropy after target recognition is *H*_*be f ore*_(Δ) − *H*_*a f ter*_(Δ) = log_2_ (*G*) − *H*(*D*) = *R*_*spacer*_ (by the definition of Rspacer in Eq. 10), and the productive interaction between the elements therefore dissipates at least ℰ_*min*_ *R*_*spacer*_ joules. Therefore, a complete target recognition event would require the dissipation of at least ℰ_*min*_ (*R*_*sequence*_ + *R*_*spacer*_) joules in total. In the pre-recruitment mechanism, the recognition complex is pre-assembled and then searches the DNA molecule as a single entity. After assembly, the recognizers in the recognition complex form a flexible structure. For any recognizer pair, the distribution of distances between the two recognizers Δ is not uniform, since it is determined by the biophysical constraints of the complex. Hence, we obtain that *H*_*be f ore*_(Δ) ≤ log_2_ (*G*), because *H*(Δ) is maximized and equal to log_2_ (*G*) only when the distribution is uniform. Therefore, *H*_*be f ore*_(Δ) − *H*_*a f ter*_(Δ) ≤ *R*_*spacer*_. The complex structure can be optimized by natural selection so that the probability distribution of the distance between the recognizers, Δ, matches the distribution *D* found in the targets. In the limit when the match between the two distributions is perfect, *H*_*be f ore*_(Δ) = *H*_*a f ter*_(Δ) = *H*(*D*), and therefore no information is gained about the distance between recognizers when target recognition occurs. Consequently, for an optimized pre-recruited recognition complex, the minimum energy dissipation during target recognition can be as low as ℰ_*min*_ *R*_*sequence*_ joules in total, regardless of the value of *R*_*spacer*_.

#### 2.3.2. Search efficiency

The difference in *H*_*be f ore*_(Δ) between recruitment and pre-recruitment mechanisms also has a significant impact on the time required for target recognition. The space of configurations for a composite motif system with n sequence patterns has *G*^*n*^ possible states, among which the recognition complex must identify a subset of *γ* targets. In the recruitment mechanism, with no a priori information on the distance between any given pair of recognizers, Δ is uniformly distributed before target recognition, *H*_*be f ore*_(Δ) = log_2_ (*G*), and the system spends an equal amount of time on all *G*^*n*^ possible states, leading to a computational search complexity of 𝒪(*G*^*n*^). In contrast, in the pre-recruitment system, complex assembly constrains the distances between recognizers and therefore reduces *H*_*be f ore*_(Δ). In a system that has been optimized by natural selection so that the distribution in recognizer distances approaches the spacer size distribution (Δ ∼ *D*), the recognition complex will spend little or no time on configurations with spacer sizes that are rare or not found among the targets, focusing instead on configurations that are frequent among the targets. For example, consider the case of a dyad-based system with a spacer of 5 bp in half of the targets and 6 bp in the other half. If the two recognizers are recruited, they have to search among *G*^2^ configurations. Instead, a pre-recruited composite recognizer can have limited flexibility, constraining the distance between the two recognizers so that only configurations where the spacer is 5 or 6 bp are explored. In this way, it can reduce the search space from *G*^2^ to 2*G*, leading to a computational search complexity of 𝒪(*G*) instead of 𝒪(*G*^2^). Hence, by focusing the search on the subset of recognizer configurations that match the targets, the pre-recruitment mechanism provides a direct advantage over recruitment in the average time required for target recognition.

### 2.4. Evolution of composite recognizers

#### 2.4.1. A Gaussian model for connectors

The operation of composite motif systems hinges on the ability of natural selection to optimize the structure of the recognition complex to closely fit the distribution of spacer sizes found in the targets. This fine-tuning ability depends on the biophysical properties that govern the interactions between recognizers, prompting the search for a simple yet realistic biophysical model capable of describing these interactions in proteins and other biological macromolecules. Formally, we define a *connector* as the model that specifies the physical distance between two sequence recognizers. In the simulations reported in Fig. 4B and 4C, we used a truncated uniform distribution for connectors, which is a simple and convenient model but not grounded in biophysics. The structure of a macromolecular complex can in principle entail very different probability distributions for the distance between two sequence recognizers, such as a bimodal distribution due to the presence of two stable conformations. However, the simplest biophysical process that can describe conformational variation in biological macromolecules is intrinsic flexibility, often mediated by the presence of particularly flexible hinge regions in proteins and protein complexes [31, 32, 33, 34]. For an intrinsically flexible protein or protein complex, we can summarize the overall stiffness of the structure with a single spring constant, *κ* [35, 36]. Thanks to this simplification, each connector in a recognition complex can be modeled as a harmonic oscillator in a thermal bath (the choice of a classical harmonic oscillator over a quantum harmonic oscillator is justified in Appendix Appendix B). The energy of each state can be found using Hamiltonian mechanics, and their probability density is obtained by applying the Boltzmann distribution (see Methods 3.2). The resulting probability density function is a Gaussian distribution with mean *µ*_*protein*_ equal to the distance when in the relaxed state, and standard deviation 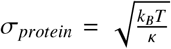. In other words, the variance of the physical distance is directly proportional to the thermal energy and inversely proportional to the stiffness of the protein structure:

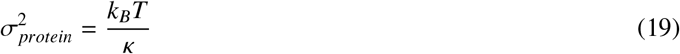

This result can also be obtained by applying the equipartition theorem (see Appendix Appendix A.1).

Eq. 19 is directly interpretable in pre-recruitment systems, where the Gaussian distribution 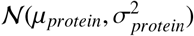 models the distance between two recognizers connected by a flexible structure. The Gaussian model is also valid for recruitment-based systems, in which each recognizer binds to the nucleotide polymer independently, and after binding, their distance is also normally distributed, with a variance that depends on the flexibilities of all the recognizers (see Appendix Appendix C).

Using the harmonic oscillator model in our simulations, we can easily keep track of the process of target search optimization as the regulator adapts to its targets. Indeed, we expect that the size distribution *D* of the targets’ spacers and the distance distribution Δ between recognizers (dictated by the spring constant *κ*) should match. If we convert the units of *D* to express a physical distance instead of using base pairs, we can define 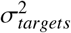 as the variance of the physical distance between the two sequence patterns, and the optimal spring constant *κ*_*opt*_ as the value of *κ* such that 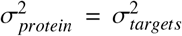. Therefore, we expect that, by natural selection, the protein complex will evolve a stiffness *κ* ≈ *κ*_*opt*_. The match is only approximate because, differently from Δ, the distribution of space sizes *D* cannot truly be Gaussian, due to the discreteness imposed by the finite value of *γ*. Moreover, the unit conversion for *D* implies a linear relationship between genomic and physical distance that does not hold for very long spacers, due to the flexibility of the DNA molecule.

#### 2.4.2. Evolution of stiffness in the recognition complex

The theory put forward in this work does not make any assumption about how the composite recognizer works. Eq. 17 is a statement about how much information should be encoded in the targets and not about the recognition mechanism. Therefore, the results hold for any models of sequence recognizers and connectors, and for any scoring function, as long as they can evolve parameters that allow them to specialize on specific patterns sufficiently. Our evolutionary simulations can run with connectors modeled as a Uniform or as a Gaussian probability density function (Methods 3.1). As expected, the results obtained with uniform connectors shown in Fig. 4B and 4C are general and can be recapitulated also with Gaussian connectors (Suppl. Fig. S1). We ran simulations for dyad-based systems using Gaussian connectors to study the evolution of the spring constant *κ*, which defines the stiffness of the recognition complex structure that recognizes the composite motif, in order to test our expectation that values of *κ* that are close to *κ*_*opt*_ should be favored. We found a strong selective pressure for the optimization of the genetically encoded stiffness of the recognition complex. Indeed, the value of *κ* becomes comparable to *κ*_*opt*_ in just a few hundred generations (Fig. 6).

**Figure 6:**
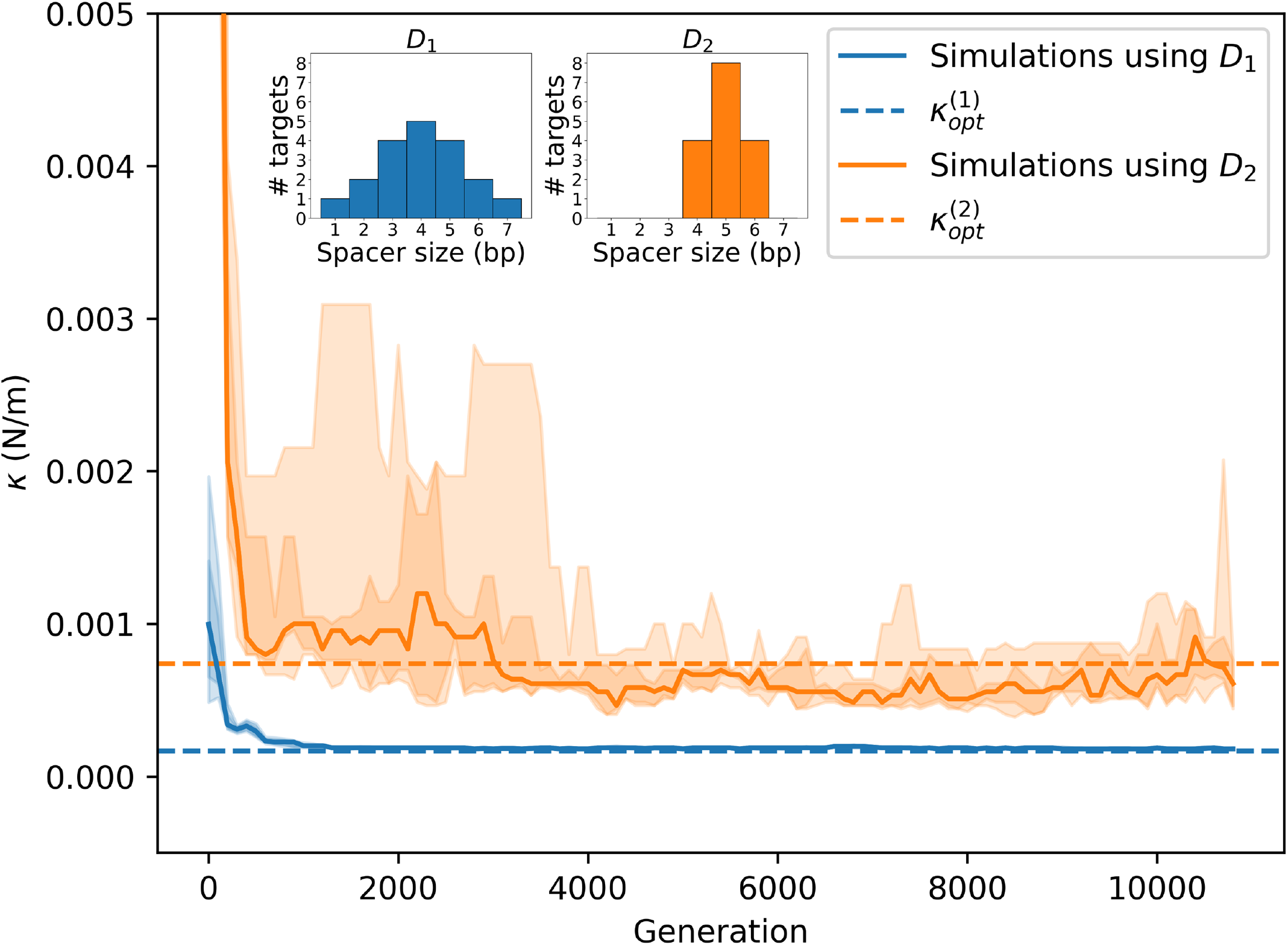
Evolution of the spring constant (*κ*) modeling the flexibility of the regulator in dyad-based systems (*n* = 2). The simulations were run using two alternative spacer size distributions *D*_1_ and *D*_2_ (shown as histograms). The plot shows the evolution of the spring constant (*κ*) of the best organism in a population of 64 organisms. 25 independent simulations were run for each of the two spacer size distributions. The solid line represents the median, and the shaded areas denote the central 50% (interquartile range) and 75% of the data. Runs using the distribution *D*_1_ (in blue) had the parameters *G* = 560, *γ* = 19. For those runs, the predicted optimal spring constant value is 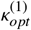. Runs using the distribution *D*_2_ (in orange) had the parameters *G* = 512, *γ* = 16. For those runs, the predicted optimal spring constant value is 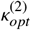. The values of 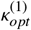 and 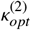 are shown as dashed lines.

The results of our simulations with the Gaussian model demonstrate that the recognition of spacer size can be based on the optimization of the stiffness of the recognition complex. To check if this recognition mechanism is compatible with the biophysics of macromolecules, we calculated the values of *κ*_*opt*_ for biologically plausible scenarios and compared them with the estimated values for proteins reported in the literature. Based on current knowledge of composite sequence motifs targeted by bacterial transcription factors [14, 15, 16, 17], we found that *κ*_*opt*_ would have to be at least 10^−2^ *N*/*m* in the case of fully conserved spacer size, and up to around 10^−4^ *N*/*m* for highly variable spacers (up to *Var*(*D*) = 4) [37]. These values are biologically plausible, as they sit right inside the range of observed values of *κ* in protein structures (i.e., from 10^−5^ to 1 N/m) [35, 38, 39, 40, 41].

### 2.5. Mutational robustness of different information-encoding strategies

According to the theory we have described so far, any system that satisfies Eq. 15 (i.e. a functional system lying on the line segment shown in Fig. 3B) can achieve perfect classification performances, identifying its target sites without errors using *R*_*sequence*_ and *R*_*spacer*_ to varying degrees. All these systems are on equal footing at the maximum “height” available in the fitness landscape, and, therefore, it is not obvious why *R*_*sequence*_ or *R*_*spacer*_ would ever be prioritized during biological evolution. However, most known multimeric transcription factors in bacteria target rigidly-spaced sequence patterns, suggesting that there is a clear bias towards prioritizing *R*_*spacer*_. We reasoned that the success of an information-encoding strategy will not only depend on the fitness of the individual, but also on the fitness of its offspring, which are subject to mutations. Thus, individuals adopting strategies that are highly sensitive to mutations can face a disadvantage, because their offspring will tend to have lower fitness compared to organisms using a strategy that is more robust to mutations. We note that nucleotide substitutions cannot change the spacer length and thus have no impact on *R*_*spacer*_. Therefore, we predict that if substitutions were the only occurring mutations, it would be advantageous to encode as much information as possible as *R*_*spacer*_, thus minimizing the value of *R*_*sequence*_ to be maintained in the face of mutations. However, real genomes are also subject to insertions and deletions (indels) that can alter spacer size, degrading *R*_*spacer*_. Therefore, we hypothesized that the optimality of the encoding strategies would be strongly dependent on the balance between the rates of these mutation types.

We validated this hypothesis by running *in silico* competition experiments between different strategists under varying mutational regimes (Fig. 7).

**Figure 7:**
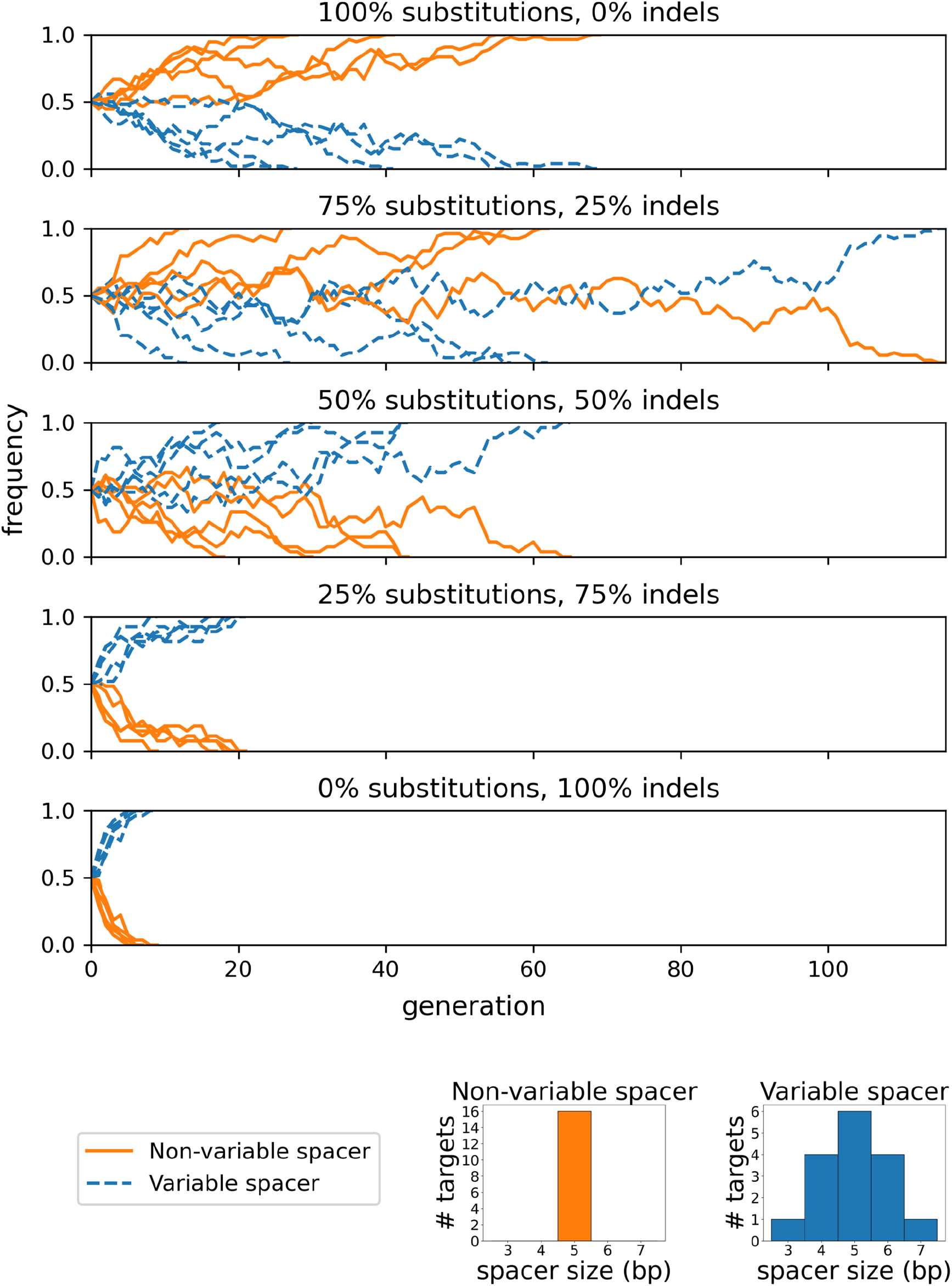
Competition experiments between evolved dyad-based systems (*n* = 2) targeting *γ* = 16 sites with variable and non-variable spacers on a genome of size *G* = 512 bp. Each panel reports 5 independent competitions under different mutational regimes. The target spacer size distributions are shown as histograms.

In each experiment, half of the organisms use a maximally conserved spacer (no spacer size variability), while the other half uses a variable spacer, with a variance that is common in reported composite transcription factor binding motifs [37]. Importantly, both competing strategies are represented by organisms that had been previously evolved to convergence, achieving perfect performances (they can all recognize their targets without errors). Thus, systematic dominance of one strategy over the other cannot be due to their performances at the start of the competition experiment, but only to the performances of their descendants, whose functionality is affected by mutations. We ran the competition experiment under five different scenarios, progressively changing the balance between substitution and indel rates. Our results (Fig. 7) confirm our intuition that when substitutions dominate, spacer conservation is favored, enabling organisms to reduce the amount of information that is sensitive to substitutions (*R*_*sequence*_) by encoding as much information in the spacer as possible. In contrast, as the relative frequency of indels increases, fixed spacer strategies become very sensitive to mutation, and it becomes advantageous to reduce spacer conservation (*R*_*spacer*_) and increase nucleotide conservation (*R*_*sequence*_) accordingly.

## 3. Methods

### 3.1. Evolutionary simulations

Each “organism” is encoded as a genome, defined as a sequence of *G* nucleotides. The first positions encode for *n* genes, each one modeling one of the *n* recognizer elements of the transcriptional regulatory complex, as well as *n* − 1 genes that encode for the biophysical properties of the joints between recognizer elements, which we call connectors (Fig. 8). The strings of nucleotides in the *n* genes for the recognizer elements are translated into *n* position-specific scoring matrices (PSSMs), the classic model for TF binding preferences. Each of those *n* genes encodes a set of *m* sequences that are aligned and used to compute nucleotide frequencies at each position of the motif (thus, *m* defines the resolution in frequency space, and its default value is 10). Instead, each of the *n* − 1 gene sequences for connectors encodes two numbers. Their role depends on the chosen model for connectors. With the *Uniform* model, the two numbers encode for the minimum and maximum spacer size allowed by the connector, and each allowed value is assigned the same probability. Thus, the two numbers represent the left (*l*) and right (*r*) bounds of a discrete uniform distribution. With the *Gaussian* model, one value represents the “relaxed state” (the mean of the Gaussian, *µ*), while the other one represents the flexibility of the protein structure (the spring constant, *κ*, based on which the standard deviation of the Gaussian, σ, is estimated, following Eq. 19). The resulting Gaussian distribution is truncated at 3 standard deviations from the mean (infinite support is unrealistic, and the harmonic oscillator model is less and less accurate as the system moves away from the relaxed state).

**Figure 8:**
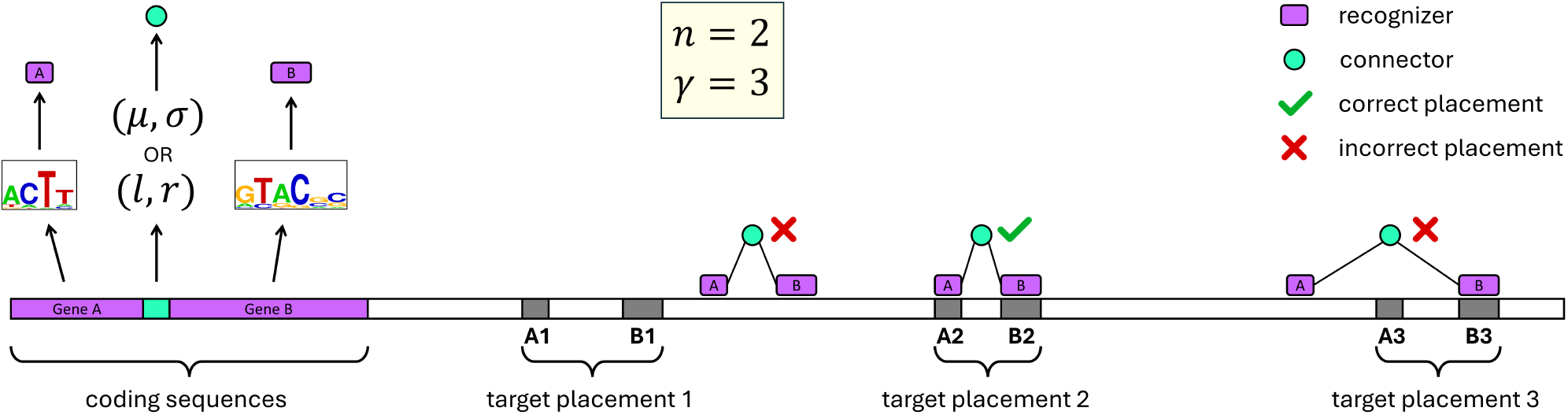
Representation of an “organism”. The genome sequence starts with the coding sequences that encode the parameters of the regulator (recognizers and connectors). The number of recognizer genes is equal to *n*, which is the specified number of patterns in the composite motif. There are *n* − 1 connector genes, each found between the genes for the two recognizers it connects. A set of *γ* target placements on the genome is randomly assigned when the organism is created. The figure represents the case where *n* = 2 and *γ* = 3.

At the beginning of a simulation, each genome in the population is initialized by randomly choosing one of the four DNA bases (each with equal probability) at each position, including the genes encoding the regulatory proteins. Then, *γ* genomic placements are randomly designated as targets in each genome. Each placement is defined by *n* genomic coordinates, each indicating the position of one of the *n* parts of the composite motif. *G, γ* and *n* are set by the user. The program scans the genome with the genetically encoded regulator, to estimate its binding affinity for each possible placement on the genome according to a scoring function (see Appendix Appendix D). The better the regulator is at distinguishing the true targets from the other placements based on binding affinity, the higher the *fitness* assigned to the organism. Specifically, the fitness function captures the classification performances of the regulator by computing the area under the precision-recall curve (AUPRC). The AUPRC ranges from 0 to 1. Perfect classification with no errors requires an AUPRC of 1. Each organism reproduces clonally, doubling the population size, and all the organisms are subject to one random mutation per generation. Then, the 50% of the population with higher fitness is selected and proceeds to the next generation. This simple selection strategy maintains the population size constant. This approach to mutation and selection was chosen due to its simplicity and because it is the one used in the original *Ev* program that was used to corroborate the classical theory [5].

As in the case of the *Ev* program, the method by which genes are translated and the scoring functions are not relevant, as long as they allow for target sites and recognizers to co-evolve to sufficient specificity. Indeed, the theory discussed here doesn’t assume any specific recognition mechanism, and it only makes statements about the overall information content of the target motifs.

The code for our evolutionary simulations is available at https://github.com/ErillLab/Info-Theo-of-Composite-Motifs.

### 3.2. Biophysical model of connectors

Modeling the flexibility of the connector with a spring constant *κ* implies that the distance between the connected recognizer elements, *δ*, assumes a certain value *µ* when the system is in the relaxed state, and that the potential energy is 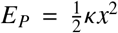, where *x* = *δ* − *µ*. Due to the size of the biological macromolecules of interest, this system can be modeled as a classical harmonic oscillator instead of a quantum harmonic oscillator (see Appendix Appendix B). Its total energy is given by the Hamiltonian *H* = *E*_*K*_ + *E*_*P*_, where 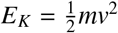 is the kinetic energy, with *m* and *v* for mass and velocity. We can express it as a function of displacement *x* and momentum *p*:

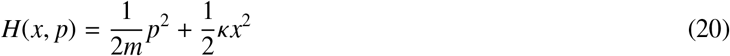

Using the Boltzmann equation, the probability density for a state with a given displacement and momentum is:

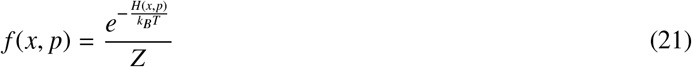

where *k*_B_ is the Boltzmann constant, *T* is the temperature, and the partition function is 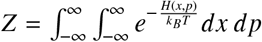. By solving the integrals, we find that *f* (*x, p*) = *f* (*x*) · *f* (*p*), which means that we can study displacement and momentum independently (see Appendix A.2). The marginal distribution for displacement is:

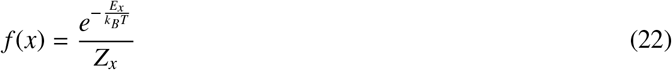

where 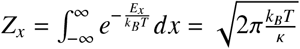. SInce *x* =*δ*− *μ*, we find that the distance *δ* follows the probability density function:

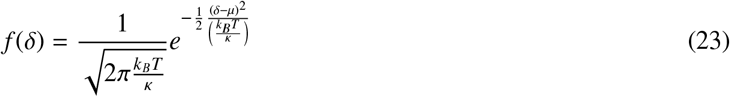

which is a Gaussian distribution with mean *µ* and standard deviation 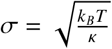.

## 4. Discussion

### 4.1. An information theoretical framework for composite sequence motifs

Sequence motifs are fundamental information-encoding strategies in biology, and their theoretical characterization over the past decades has fostered the development of bioinformatics algorithms to analyze transcriptional regulatory networks and other biological systems [42, 43, 6, 44, 45]. In this work, we generalize the theoretical framework of sequence motifs to characterize *composite* sequence motifs, which are composed of a series of sequence patterns separated by spacers whose length can be variable. This generalization entails the redefinition of the well-established *R*_*frequency*_ and *R*_*sequence*_ information metrics, and the introduction of *R*_*spacer*_, a new metric that quantifies the information encoded as the conservation of the size of the spacers between sequence patterns. We show that *R*_*spacer*_ can be interchanged with *R*_*sequence*_ (within certain bounds that we derive) as long as their sum equals *R*_*frequency*_. Importantly, when there are no spacers or when the spacers are not variable, our formalism reduces to the one presented in the classic theory developed for simple sequence motifs. The classical information theory framework for sequence motifs was not originally developed to model composite sequence motifs. Indeed in [1] the authors note that in the presence of variable-length spacers, *R*_*sequence*_ can be greater than the value of *R*_*frequency*_ that they defined, which is − log_2_ (*G*). Intuiting that the uncertainty in the spacer size needs to be taken into account, in the Methods section they suggest a correction term for their equation that would make the near equivalence work even if there is a variable-length spacer. This correction was introduced without a formal justification, but it was later effectively applied to model the composite motifs defining ribosomal binding sites and splice sites [6, 45]. Our generalized framework readily explains why the suggested correction works and how to generalize it (see Appendix Appendix E). Our work hence provides a unified framework to study composite and classical motifs, enabling us to investigate different aspects of the evolvability and efficiency of biological systems based on the recognition of composite motifs.

### 4.2. Efficiency considerations for recruitment and pre-recruitment strategies

Our formal analysis of composite sequence motif systems enables us to compare alternative strategies for target recognition by regulatory complexes. Both in prokaryotes and eukaryotes, biological processes like transcriptional regulation or translation initiation require the recognition of target sites, which can be accomplished through the binding of a preformed complex (pre-recruitment) or through the gradual assembly of the complex on nucleotide sequences (recruitment). Our results show that a pre-recruited recognition complex can be optimized so that the distribution of the distance between its constituent recognizers approximates the distribution of the size of the corresponding spacers in the nucleotide sequence (Fig. 6). This match in recognizer distance and spacer size distributions means that a pre-recruited complex will explore almost exclusively configurations with spacer sizes matching its target sites, decreasing the time required to locate them. Furthermore, this match in distributions means that in a pre-recruited system the minimal energy dissipation per recognition event is ℰ_*min*_ · *R*_*sequence*_ joules, instead of the ℰ_*min*_ · (*R*_*sequence*_ + *R*_*spacer*_) joules required by a recruitment mechanism.

Establishing the minimum energy dissipation for the binding of a recognition complex to a nucleotide sequence can provide a means to predict the actual amount of energy that is dissipated per recognition event. Indeed, bistate molecular machines were found to work optimally at about 70% isothermal efficiency, because an isothermal efficiency of *ln*(2) ≈ 0.69 is the highest that guarantees to operate with as few errors as desired when the effect of thermal noise is taken into account (i.e., without exceeding channel capacity) [46]. With this insight, it is possible to turn the lower bound on energy dissipation into a prediction of the amount of energy dissipated during molecular machines’ operations, because ℰ_*min*_/E = 0.7, where ℰ is the actual energy dissipated per gained bit. Predictions obtained in this way were experimentally validated for a diverse set of molecular machines, including sequence motif recognizers like transcription factors and restriction enzymes [29]. Hence, one can predict that the energy dissipated per recognized target is approximately ℰ_*min*_ · (*R*_*sequence*_ + *R*_*spacer*_) /0.7 joules for a recruitment-based system and ℰ_*min*_ · *R*_*sequence*_/0.7 joules for a pre-recruitment-based system.

The energetic benefits of a pre-recruitment strategy extend beyond the reduced energy dissipation per binding event discussed above. In the recruitment mechanism, recognition of target sites is accomplished through the sequential binding of *n* independent recognizers to their cognate binding sites, eventually resulting in the formation of a productive recognition complex. Before the recognition complex is fully assembled, thermal fluctuations may result in the disengagement of any of the recognizers. This implies that the total number of binding events required for the full assembly of the recognition complex may be larger than *n*, increasing the time and energy dissipation required for target recognition. The presence of unproductive binding events in the recruitment mechanism can be minimized by a large enough concentration of recognizers. In contrast, in the pre-recruitment mechanism, recognition of the *n* sequence patterns defining the composite motif takes place simultaneously, limiting the number of sequence binding events per target recognition to *n*, and allowing pre-recruitment systems to operate at lower recognizer concentrations.

These results raise the question of why recruitment processes would be used at all in biological systems. Notwithstanding its energetic and search efficiency limitations, the recruitment mechanism presents notable advantages. Instead of a single, preassembled recognition complex with an essentially binary output across all targets, recruitment systems may evolve to enable the formation of multiple partial complexes, which can have different regulatory activity on each target. This strategy exploits the combinatorial power of the composite motif (up to 2^*n*^ output states) and is a hallmark of transcriptional regulation in eukaryotes [8, 47, 48]. In addition, the recruitment mechanism enables the sharing of recognizers across regulatory systems, maximizing the availability of recognizers that are shared across concurrent transcriptional programs. Rather than being preemptively sequestered in one or more pre-assembled complexes, in a recruitment system a shared recognizer may be recruited to different composite motifs in conjunction with a variety of other recognizers to instantiate different regulatory responses [47, 48].

The different advantages of recruitment and pre-recruitment strategies suggest that pre-recruitment systems should be primarily exploited when there is no regulatory need to assemble partial recognition complexes and when the recognition complex does not integrate components that are shared by concurrent regulatory programs. Indeed, cells employ a mixture of recruitment and pre-recruitment strategies operating on this basic principle. In bacteria, most transcription factors are pre-recruited multimeric complexes that respond to an individual signal with an essentially binary output. These transcription factors are independently recruited to promoters, where they interact with RNA polymerase and other transcription factors for the formation of an open transcription complex. This approach exploits the combinatorial power of the recruitment mechanism for signal integration by a set of transcription factors, while at the same time maximizing the shared availability of the RNA polymerase holoenzyme across concurrent regulatory programs. Bacteria, however, also implement mutually exclusive regulatory programs through the pre-recruitment of sigma factors by the bacterial core RNA polymerase [49]. The newly formed RNA polymerase complexes not only define a new transcriptional program, but also deplete the availability of RNA polymerase complexes capable of binding promoters targeted by other sigma factors, prompting a global transition to a new cell state [50, 51, 52]. Prerecruitment of a shared recognizer may also be promoted to enact fast responses, such as in the response to superoxide in *E. coli*, in which the SoxS transcription factor is pre-recruited by RNA polymerase [53].

### 4.3. Impact of mutation rates on the prevalence of rigid spacers

The theoretical framework for composite sequence motifs developed in this work establishes that the information encoded in *γ* target sites can be expressed as the sum of the information encoded in the *n* sequence patterns (*R*_*sequence*_) and in the size distribution of the *n* − 1 spacers separating them (*R*_*spacer*_). For a composite motif-based system to be functional, this information must be approximately equal to the information (*R*_*f requqnecy*_) required to locate *γ* target sites in a genome of size *G* (Eq. 17). Hence, whenever the sum *R*_*sequence*_ + *R*_*spacer*_ satisfies this condition, information can be freely distributed among these two quantities, restricted only by the bounds that we derived (Eq. 12 and Eq. 14).

Building on the evolutionary simulation framework used to validate the near equality of *R*_*sequence*_ and *R*_*f requqnecy*_ in classical motifs, we show that this basic principle extends to composite motifs, and that functional composite motif-based systems can be evolved across the entire range of *R*_*sequence*_ and *R*_*spacer*_ value combinations that satisfy Eq. 17 and the informational bounds for *R*_*spacer*_ (Fig. 4). This observation raises an intriguing question: if composite motif-based systems can in principle adopt any of these *R*_*sequence*_ and *R*_*spacer*_ value combinations, why do most known bacterial multimeric transcription factors target composite motifs with non-variable spacers? A plausible explanation might be that the recognition of composite motifs with variable spacers requires complex conformational changes in the recognition complex, and that the evolution of such complexity is unwarranted, given that the same functional task can be accomplished with a non-variable spacer motif. However, our work shows that the intrinsic flexibility of protein structures is sufficient to enable binding to the moderately variable spacers of currently known composite motifs, and that adequate spring constant values will readily evolve to match variable-spaced composite dyad motifs in evolutionary simulations (Fig. 6).

In the absence of biochemical or structural limitations on the ability of the recognition complex to recognize composite motifs with variable spacers, an alternative explanation is that bacterial composite motifs with non-variable spacers provide an evolutionary advantage based on the way information is encoded in the genome. Different mutation types have fundamentally different effects on the different sources of information present in composite sequence motifs (*R*_*sequence*_ and *R*_*spacer*_). While the information in sequence patterns (*R*_*sequence*_) is vulnerable to both substitutions and indels, the information encoded by the size of the spacer separating sequence patterns (*R*_*spacer*_) is only sensitive to indels. This asymmetry between the two information sources of a composite sequence motif therefore provides a putative causal explanation for the prevalence of composite motifs with rigid spacers. To address this question, we performed competition experiments between pre-evolved composite motif-based systems targeting a dyad motif with a variable and a non-variable spacer, exposing them to varying ratios of substitution and indel mutation rates. Our results (Fig. 7) reveal that when substitutions dominate, non-variable spacer strategies achieve fixation in the population. With few or no indels disrupting the spacer, using a non-variable spacer to store the maximum possible amount of information in *R*_*spacer*_ maximizes mutational robustness. As the proportion of indels increases, however, spacer size variability becomes increasingly favored, and systems based on variable spacers end up systematically dominating the population (Fig. 7).

The range of indel-to-substitution ratios that favors the evolution of non-variable spacers is in agreement with the experimentally determined ratios between substitution and indel rates in bacteria, which range from 1:3 in *Bacillus subtilis* to 1:10 in *Agrobacterium tumefaciens* [54]. This aligns with the observation that most multimeric transcription factors in bacteria target non-variable spacer composite motifs [14] and suggests that bacterial transcription factors have evolved to adopt information-encoding strategies that are resilient in their mutational environment. The dominance of non-variable spacer composite motifs can be further explained by the sequence-dependent nature of indels [55, 56, 57, 58], which is heavily determined by the presence of homopolymers and simple repeats [55, 56] and which therefore makes it possible to evolve target sequences that minimize the probability of indels disturbing the spacer. Eukaryotic organisms present similar indel-to-substitution ratios [54] and, indeed, many mono- and heterodimeric transcription factors in eukaryotes target non-variable spacer composite motifs [59]. Beyond multimeric transcription factors, which typically integrate a small number of tightly coupled signals, we argue that the need to implement target-specific programs through the recruitment of a diverse set of recognition complexes can overcome the mutational advantage of non-variable spacer strategies, leading to the preponderance of flexible spacer composite motifs seen in eukaryotic regulatory modules [11].

## 5. Acknowledgments

Elia Mascolo was funded by the Merck Data Science for Observational Research Program through the UMBC College of Natural and Mathematical Sciences Academic Fellowship Program. Evolutionary simulations were run using the UMBC High Performance Computing Facility (HPCF). The facility is supported by the US National Science Foundation through the MRI program (CNS-0821258, CNS-1228778, OAC-1726023 and CNS-1920079) and the SCREMS program (DMS-08215311), with additional support from the University of Maryland, Baltimore County (UMBC).

## Appendix A. Relationship between *Var*(Δ) and *κ*

### Appendix A.1. Using the equipartition theorem

The equipartition theorem states that, in thermal equilibrium, each degree of freedom is associated with a mean energy of 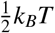 joules [60]. For the one-dimensional harmonic oscillator in a thermal bath we have that the average potential energy and kynetic energy are 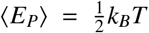 and 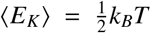, respectively. The use of the classical harmonic oscillator and the use of the equipartition theorem are justified in Appendix B. By combining it with the definition 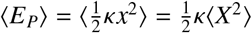, we find

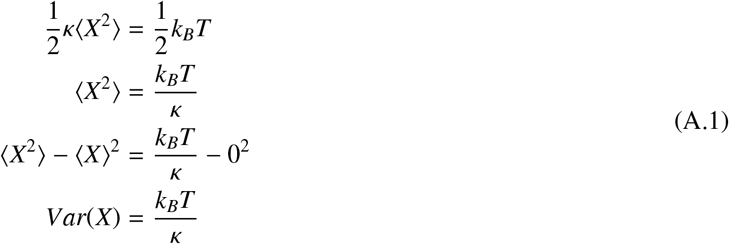

Since *x* = *δ* − *µ*, we have that *X* is just a translation of Δ such that *X* = Δ − *µ*, and therefore *Var*(Δ) = *Var*(*X*). Thus, also 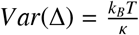.

### Appendix A.2. Independence of displacement and momentum

In Eq. 21, the partition function is 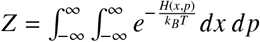. Thus,

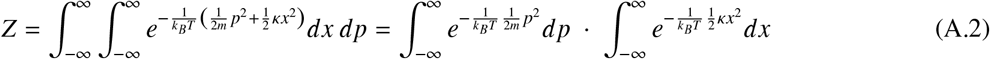

By solving the two Gaussian integrals, we find 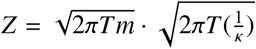, which is the product of two terms such that the first term is only a function of momentum *p*, while the second term is only a function of displacement *x*. We can call them the marginals *Z*_*p*_ and *Z*_*x*_, respectively. So, *Z* = *Z*_*p*_ · *Z*_*x*_, and Eq. 21 can be re-written as

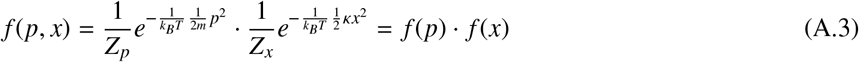

Thus, the joint probability density is simply the product of the marginal probability densities *f* (*p*) and *f* (*x*). Hence, momentum and displacement are independent.

## Appendix B. Classical harmonic oscillator

The fact that the energy levels are discrete can require modeling oscillators as quantum harmonic oscillators. The classical harmonic oscillator is a good approximation only if the energy step is much smaller than the average thermal energy, i.e., when *h*ν ≪ *k*_*B*_*T*, where *h* is the Planck’s constant and 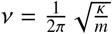 is the classical frequency of the oscillator, with *m* being the mass and *κ* the spring constant. So, the discreteness of the energy levels is negligible when 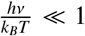. We considered an extremely comprehensive range of possible spring constant values for protein structures obtained from the literature [35, 38, 39, 40, 41], namely the range [10^−5^, 1] (newtons per meter). In terms of mass, we considered a range from just 20 kDa (3.3 · 10^−23^ kg) for very small transcription factors up to 200 kDa (3.3 · 10^−22^ kg) for very large ones. Finally, we considered that the temperature inside a living cell could range from around −20 ^°^*C* (253.15 K) in some psychrophiles up to around 110 ^°^*C* (383.15 K) in some thermophiles. With these ranges, we found that 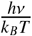 can have values that range from 10^−13^ up to only 10^−10^. Therefore, we can say that 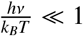, and that the use of a classical harmonic oscillator is justified, as well as the use of the equipartition theorem.

## Appendix C. Generality of Gaussian connector models

We have shown that for a *pre-recruited* complex with some flexibility modeled with a spring constant (*κ*), the distance between sequence recognizers (like two DNA binding domains of a TF complex) is normally distributed. Here we show that the same holds for *recruited* complexes, in which the components bind sequences independently, and they do or do not form a complex only after they are bound.

Consider two recognizers already bound to two sequence segments. They could differ in terms of flexibility (as in the case of a heterodimeric complex), therefore we name their spring constants *κ*_1_ and *κ*_2_. We have already noted that their position coordinates are normally distributed along any direction. The distance between the two recognizers is 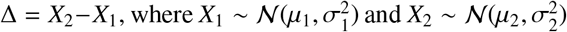 are their positions along the direction determined by the vector joining their positions, which is assumed to be roughly parallel to the nucleotide sequence on which they are placed. We also know that 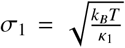 and 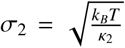 (see Methods 3.2). Thus, the distance between recognizers is also normally distributed, with a mean distance *µ*_*δ*_ = *E*[Δ] = *µ*_2_ −*µ*_1_ and variance 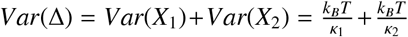. We can re-write the variance of the distance as 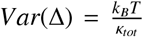, where 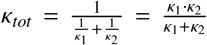, i.e., twice the harmonic mean between *κ*_1_ and *κ*_2_. The standard deviation 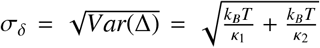 is a function of the flexibilities of both the recognizer structures, as expected. Therefore, the use of a Gaussian connector to model protein complex flexibility is valid not only for systems based on pre-recruitment, but also for those based on recruitment, where we have that 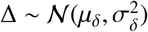.

## Appendix D. Scoring function for composite recognizers used in the evolutionary simulations

Composite recognizers, on top of being influenced by the sequence preferences of each sequence recognizer, also have preferences for the distance (in bp) between the placements of the sequence recognizers. Thus, the score of each placement must be a function of both the observed sequences and the distance between them. The conventional approach to modeling sequence specificity is the position-specific scoring matrix (PSSM) [19]. The scoring function used when scanning sequences with PSSMs is a log-likelihood ratio. This approach is widely used because the likelihood ratio is a uniformly most powerful statistical test, according to the Neyman–Pearson lemma [61]. Therefore, in the absence of specific priors, the likelihood ratio (and thus also the log-likelihood ratio) ranks observed sequences in a way that maximizes classification performances. We extended the log-likelihood ratio framework to include all the observations made by composite recognizers, i.e., *n* sequences and *n* − 1 spacer lengths. The null hypothesis *H*_0_ states that the observation is due to chance, while the alternative hypothesis *H*_1_ states that it is sampled from the set of true targets. The test assumes that the different observations are independent events (or that their correlations are negligible). For example, for *n* = 2 each placement is scored as the log-likelihood ratio

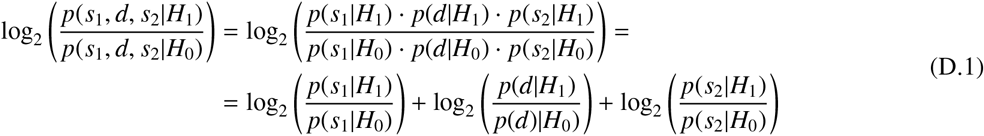

where *s*_1_, *d, s*_2_ are the observed features of the placement, i.e., the first sequence pattern, the spacer size and the second sequence pattern, respectively. This equivalence shows that the scoring function is a log-likelihood ratio that can be expressed as the sum of the conventional PSSM scores plus the connector scores, each being a log-likelihood ratio itself. The null hypothesis is tested using a background model. For PSSMs, the background model assumes that at each genomic position, the probability of each nucleotide is equal to its frequency in the genome (a 0-th order Markov model), while for connectors, the background model assumes all possible distance values are equally probable, with probability 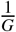 (because, on a circular genome, there are *G* pairs of positions with distance d out of *G*^2^ pairs, for each value of *d*). The alternative hypothesis is tested using the probability distributions defined by PSSMs and connectors. The probability distribution of a connector can be a (truncated) uniform or Gaussian distribution. The probability *p*(*d*|*H*_1_) is obtained by integrating the probability density function specified by the connector from *d* − 0.5 to *d* + 0.5.

## Appendix E. Validity of previously proposed correction for dyad motifs

In [1], the authors address the case in which a sequence motif contains a variable-length spacer. This is what we would call a dyad motif (*n* = 2). In that case, they note that 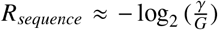 does not hold. To obtain the predicted value of 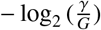 bits, they suggest a correction that consists of subtracting from *R*_*sequence*_ the uncertainty of the spacer, which we named *H*(*D*). We can write their approach as 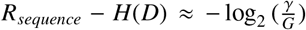. Our theory simply equates the total information from all the sources to 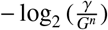 and it can be used to prove the general validity of the correction described above as follows:

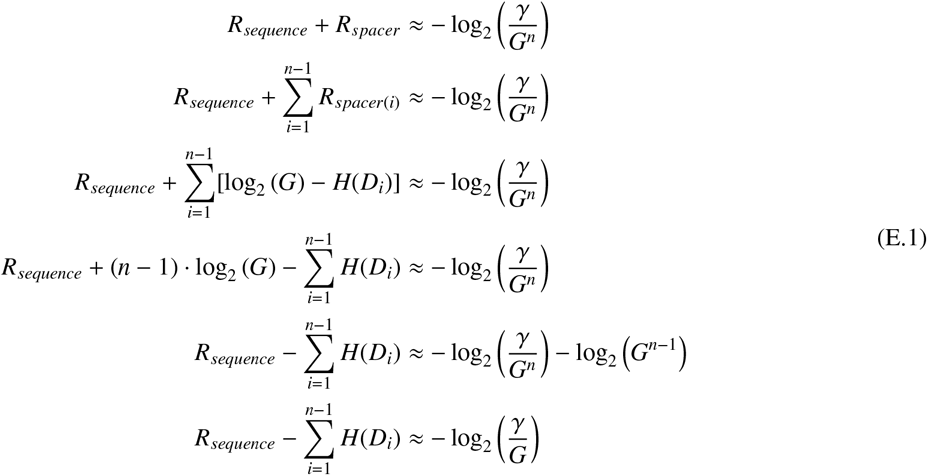

Here *H*(*D*_*i*_) is the uncertainty of the *i*-th spacer. Therefore, the uncertainty values from all the spacers need to be subtracted from *R*_*sequence*_ for the result to be 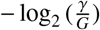. In the special case when *n* = 2, there is only one spacer, and we obtain the correction proposed by Schneider *et al*.

## References

[1] T. D. Schneider, G. D. Stormo, L. Gold, A. Ehrenfeucht, Information content of binding sites on nucleotide sequences, Journal of Molecular Biology 188 (3) (1986) 415–431. doi:10.1016/0022-2836(86)90165-8. URL https://www.sciencedirect.com/science/article/pii/0022283686901658

[2] C. D. Manning, P. Raghavan, H. Schütze, Introduction to Information Retrieval, ISBN: 9780511809071 Publisher: Cambridge University Press (Jul. 2008). doi:10.1017/CBO9780511809071. URL https://www.cambridge.org/highereducation/books/introduction-to-information-retrieval/669D108D20F556C5C30957D63B5AB65

[3] G. E. Crooks, G. Hon, J.-M. Chandonia, S. E. Brenner, WebLogo: A Sequence Logo Generator, Genome Research 14 (6) (2004) 1188–1190, company: Cold Spring Harbor Laboratory Press Distributor: Cold Spring Harbor Laboratory Press Institution: Cold Spring Harbor Laboratory Press Label: Cold Spring Harbor Laboratory Press Publisher: Cold Spring Harbor Lab. doi:10.1101/gr.849004. URL https://genome.cshlp.org/content/14/6/1188

[4] C. E. Shannon, A mathematical theory of communication, The Bell System Technical Journal 27 (3) (1948) 379–423, conference Name: The Bell System Technical Journal. doi:10.1002/j.1538-7305.1948.tb01338.x. URL https://ieeexplore.ieee.org/document/6773024

[5] T. D. Schneider, Evolution of biological information, Nucleic Acids Research 28 (14) (2000) 2794–2799. doi:10.1093/nar/28.14.2794. URL https://doi.org/10.1093/nar/28.14.2794

[6] R. K. Shultzaberger, R. E. Bucheimer, K. E. Rudd, T. D. Schneider, Anatomy of Escherichia coli ribosome binding sites1, Journal of Molecular Biology 313 (1) (2001) 215–228. doi:10.1006/jmbi.2001.5040. URL https://www.sciencedirect.com/science/article/pii/S0022283601950405

[7] L. Wachutka, L. Caizzi, J. Gagneur, P. Cramer, Global donor and acceptor splicing site kinetics in human cells, eLife 8 (2019) e45056, publisher: eLife Sciences Publications, Ltd. doi:10.7554/eLife.45056. URL https://doi.org/10.7554/eLife.45056

[8] A. Reményi, H. R. Schöler, M. Wilmanns, Combinatorial control of gene expression, Nature Structural & Molecular Biology 11 (9) (2004) 812–815, publisher: Nature Publishing Group. doi:10.1038/nsmb820. URL https://www.nature.com/articles/nsmb820

[9] I. Georgakopoulos-Soares, C. Deng, V. Agarwal, C. S. Y. Chan, J. Zhao, F. Inoue, N. Ahituv, Transcription factor binding site orientation and order are major drivers of gene regulatory activity, Nature Communications 14 (1) (2023) 2333, publisher: Nature Publishing Group. doi:10.1038/s41467-023-37960-5. URL https://www.nature.com/articles/s41467-023-37960-5

[10] S. H. Duttke, C. Guzman, M. Chang, N. P. Delos Santos, B. R. McDonald, J. Xie, A. F. Carlin, S. Heinz, C. Benner, Position-dependent function of human sequence-specific transcription factors, Nature 631 (8022) (2024) 891–898, publisher: Nature Publishing Group. doi:10.1038/s41586-024-07662-z. URL https://www.nature.com/articles/s41586-024-07662-z

[11] V. J. Makeev, A. P. Lifanov, A. G. Nazina, D. A. Papatsenko, Distance preferences in the arrangement of binding motifs and hierarchical levels in organization of transcription regulatory information, Nucleic Acids Research 31 (20) (2003) 6016–6026. doi:10.1093/nar/gkg799. URL https://doi.org/10.1093/nar/gkg799

[12] C. K. Glass, Differential Recognition of Target Genes by Nuclear Receptor Monomers, Dimers, and Heterodimers*, Endocrine Reviews 15 (3) (1994) 391–407. doi:10.1210/edrv-15-3-391. URL https://doi.org/10.1210/edrv-15-3-391

[13] D. Panne, The enhanceosome, Current Opinion in Structural Biology 18 (2) (2008) 236–242. doi:10.1016/j.sbi.2007.12.002. URL https://www.sciencedirect.com/science/article/pii/S0959440X07002023

[14] C. Laguri, M. K. Phillips-Jones, M. P. Williamson, Solution structure and DNA binding of the effector domain from the global regulator PrrA (RegA) from Rhodobacter sphaeroides: insights into DNA binding specificity, Nucleic Acids Research 31 (23) (2003) 6778–6787. doi:10.1093/nar/gkg891. URL https://doi.org/10.1093/nar/gkg891

[15] H. Chao, N.-Y. Zhou, GenR, an IclR-Type Regulator, Activates and Represses the Transcription of gen Genes Involved in 3-Hydroxybenzoate and Gentisate Catabolism in Corynebacterium glutamicum, Journal of Bacteriology 195 (7) (2013) 1598–1609, publisher: American Society for Microbiology. doi:10.1128/jb.02216-12. URL https://journals.asm.org/doi/10.1128/jb.02216-12

[16] A. Tramonti, C. Nardella, M. L. di Salvo, S. Pascarella, R. Contestabile, The MocR-like transcription factors: pyridoxal 5-phosphate-dependent regulators of bacterial metabolism, The FEBS Journal 285 (21) (2018) 3925–3944, eprint: https://onlinelibrary.wiley.com/doi/pdf/10.1111/febs.14599. doi:10.1111/febs.14599. URL https://onlinelibrary.wiley.com/doi/abs/10.1111/febs.14599

[17] N. V. Sernova, M. S. Gelfand, Comparative Genomics of CytR, an Unusual Member of the LacI Family of Transcription Factors, PLOS ONE 7 (9) (2012) e44194, publisher: Public Library of Science. doi:10.1371/journal.pone.0044194. URL https://journals.plos.org/plosone/article?id=10.1371/journal.pone.0044194

[18] G. D. Stormo, Modeling the specificity of protein-DNA interactions, Quantitative Biology 1 (2) (2013) 115–130, eprint: https://onlinelibrary.wiley.com/doi/pdf/10.1007/s40484-013-0012-4. doi:10.1007/s40484-013-0012-4. URL https://onlinelibrary.wiley.com/doi/abs/10.1007/s40484-013-0012-4

[19] G. Z. Hertz, G. D. Stormo, Identifying DNA and protein patterns with statistically significant alignments of multiple sequences., Bioinformatics 15 (7) (1999) 563–577. doi:10.1093/bioinformatics/15.7.563. URL https://doi.org/10.1093/bioinformatics/15.7.563

[20] T. D. Schneider, Theory of molecular machines. I. Channel capacity of molecular machines, Journal of Theoretical Biology 148 (1) (1991) 83–123. doi:10.1016/S0022-5193(05)80466-7. URL https://www.sciencedirect.com/science/article/pii/S0022519305804667

[21] T. D. Schneider, Theory of molecular machines. II. Energy dissipation from molecular machines, Journal of Theoretical Biology 148 (1) (1991) 125–137. doi:10.1016/S0022-5193(05)80467-9. URL https://www.sciencedirect.com/science/article/pii/S0022519305804679

[22] J. H. Felker, A Link Between Information and Energy, Proceedings of the IRE 40 (6) (1952) 728–729, conference Name: Proceedings of the IRE. doi:10.1109/JRPROC.1952.274070. URL https://ieeexplore.ieee.org/document/4051030

[23] F. P. Adler, Comments on “{Figure of Merit for Communication Devices}”, Proceedings of the IRE 42 (7) (1954) 1188–1193, conference Name: Proceedings of the IRE. doi:10.1109/JRPROC.1954.274556. URL https://ieeexplore.ieee.org/document/4051768/?arnumber=4051768

[24] J. Von Neumann, Theory of Self-Reproducing Automata; Fourth University of Illinois lecture, burks. a. w. Edition, University of Illinois Press, Urbana, 1966.

[25] R. Landauer, Irreversibility and Heat Generation in the Computing Process, IBM Journal of Research and Development 5 (3) (1961) 183–191, conference Name: IBM Journal of Research and Development. doi:10.1147/rd.53.0183. URL https://ieeexplore.ieee.org/document/5392446

[26] J. R. Pierce, C. C. Cutler, Interplanetary Communications, in: F. I. Ordway (Ed.), Advances in Space Science, Academic Press, 1959, pp. 55–109. doi:10.1016/B978-1-4831-9959-7.50006-2. URL https://www.sciencedirect.com/science/article/pii/B9781483199597500062

[27] R. W. Keyes, R. Landauer, Minimal Energy Dissipation in Logic, IBM Journal of Research and Development 14 (2) (1970) 152–157, conference Name: IBM Journal of Research and Development. doi:10.1147/rd.142.0152. URL https://ieeexplore.ieee.org/document/5391667

[28] R. W. Keyes, Power Dissipation in Information Processing, Science 168 (3933) (1970) 796–801, publisher: American Association for the Advancement of Science. doi:10.1126/science.168.3933.796. URL https://www.science.org/doi/10.1126/science.168.3933.796

[29] T. D. Schneider, Generalizing the isothermal efficiency by using Gaussian distributions, PLOS ONE 18 (1) (2023) e0279758, publisher: Public Library of Science. doi:10.1371/journal.pone.0279758. URL https://journals.plos.org/plosone/article?id=10.1371/journal.pone.0279758

[30] T. D. Schneider, A brief review of molecular information theory, Nano Communication Networks 1 (3) (2010) 173–180. doi:10.1016/j.nancom.2010.09.002. URL https://www.sciencedirect.com/science/article/pii/S1878778910000359

[31] M. Lüking, J. Elf, Y. Levy, Conformational Change of Transcription Factors from Search to Specific Binding: A lac Repressor Case Study, The Journal of Physical Chemistry B 126 (48) (2022) 9971–9984, publisher: American Chemical Society. doi:10.1021/acs.jpcb.2c05006. URL https://doi.org/10.1021/acs.jpcb.2c05006

[32] A. Haelens, T. Tanner, S. Denayer, L. Callewaert, F. Claessens, The Hinge Region Regulates DNA Binding, Nuclear Translocation, and Transactivation of the Androgen Receptor, Cancer Research 67 (9) (2007) 4514–4523. doi:10.1158/0008-5472.CAN-06-1701. URL https://doi.org/10.1158/0008-5472.CAN-06-1701

[33] H. Sewell, T. Tanaka, K. E. Omari, E. J. Mancini, A. Cruz, N. Fernandez-Fuentes, J. Chambers, T. H. Rabbitts, Conformational flexibility of the oncogenic protein LMO2 primes the formation of the multi-protein transcription complex, Scientific Reports 4 (1) (2014) 3643, publisher: Nature Publishing Group. doi:10.1038/srep03643. URL https://www.nature.com/articles/srep03643

[34] S. C. Flores, L. J. Lu, J. Yang, N. Carriero, M. B. Gerstein, Hinge Atlas: relating protein sequence to sites of structural flexibility, BMC Bioinformatics 8 (1) (2007) 167. doi:10.1186/1471-2105-8-167. URL https://doi.org/10.1186/1471-2105-8-167

[35] R. Sarkar, Native flexibility of structurally homologous proteins: insights from anisotropic network model, BMC Biophysics 10 (1) (2017) 1. doi:10.1186/s13628-017-0034-9. URL https://doi.org/10.1186/s13628-017-0034-9

[36] J. M. McBride, J.-P. Eckmann, T. Tlusty, General Theory of Specific Binding: Insights from a Genetic-Mechano-Chemical Protein Model, Molecular Biology and Evolution 39 (11) (2022) msac217. doi:10.1093/molbev/msac217. URL https://doi.org/10.1093/molbev/msac217

[37] P. S. Novichkov, A. E. Kazakov, D. A. Ravcheev, S. A. Leyn, G. Y. Kovaleva, R. A. Sutormin, M. D. Kazanov, W. Riehl, A. P. Arkin, I. Dubchak, D. A. Rodionov, RegPrecise 3.0 – A resource for genome-scale exploration of transcriptional regulation in bacteria, BMC Genomics 14 (1) (2013) 745. doi:10.1186/1471-2164-14-745. URL https://doi.org/10.1186/1471-2164-14-745

[38] G. Zaccai, How Soft Is a Protein? A Protein Dynamics Force Constant Measured by Neutron Scattering, Science 288 (5471) (2000) 1604–1607, publisher: American Association for the Advancement of Science. doi:10.1126/science.288.5471.1604. URL https://www.science.org/doi/10.1126/science.288.5471.1604

[39] G. Haran, H. Mazal, How fast are the motions of tertiary-structure elements in proteins?, The Journal of Chemical Physics 153 (13) (2020) 130902. doi:10.1063/5.0024972. URL https://doi.org/10.1063/5.0024972

[40] A. R. Strom, R. J. Biggs, E. J. Banigan, X. Wang, K. Chiu, C. Herman, J. Collado, F. Yue, J. C. Ritland Politz, L. J. Tait, D. Scalzo, A. Telling, M. Groudine, C. P. Brangwynne, J. F. Marko, A. D. Stephens, HP1 is a chromatin crosslinker that controls nuclear and mitotic chromosome mechanics, eLife 10 (2021) e63972, publisher: eLife Sciences Publications, Ltd. doi:10.7554/eLife.63972. URL https://doi.org/10.7554/eLife.63972

[41] E. H. Lee, J. Hsin, O. Mayans, K. Schulten, Secondary and Tertiary Structure Elasticity of Titin Z1Z2 and a Titin Chain Model, Biophysical Journal 93 (5) (2007) 1719–1735. doi:10.1529/biophysj.107.105528. URL https://www.sciencedirect.com/science/article/pii/S0006349507714285

[42] S. Kılıç, M. Sánchez-Osuna, A. Collado-Padilla, J. Barbé, I. Erill, Flexible comparative genomics of prokaryotic transcriptional regulatory networks, BMC Genomics 21 (Suppl 5) (2020) 466. doi:10.1186/s12864-020-06838-x. URL https://pmc.ncbi.nlm.nih.gov/articles/PMC7739468/

[43] P. K. Rogan, B. M. Faux, T. D. Schneider,

[44] T. D. Schneider, Strong minor groove base conservation in sequence logos implies DNA distortion or base flipping during replication and transcription initiation, Nucleic Acids Research 29 (23) (2001) 4881–4891. doi:10.1093/nar/29.23.4881. URL https://doi.org/10.1093/nar/29.23.4881

[45] P. K. Rogan, Ab initio exon definition using an information theory-based approach, in: 2009 43rd Annual Conference on Information Sciences and Systems, 2009, pp. 847–852. doi:10.1109/CISS.2009.5054835. URL https://ieeexplore.ieee.org/document/5054835

[46] T. D. Schneider, 70% efficiency of bistate molecular machines explained by information theory, high dimensional geometry and evolutionary convergence, Nucleic Acids Research 38 (18) (2010) 5995–6006. doi:10.1093/nar/gkq389. URL https://doi.org/10.1093/nar/gkq389

[47] F. Reiter, S. Wienerroither, A. Stark, Combinatorial function of transcription factors and cofactors, Current Opinion in Genetics & Development 43 (2017) 73–81. doi:10.1016/j.gde.2016.12.007. URL https://www.sciencedirect.com/science/article/pii/S0959437X17300059

[48] M. Ptashne, Regulation of transcription: from lambda to eukaryotes, Trends in Biochemical Sciences 30 (6) (2005) 275–279. doi:10.1016/j.tibs.2005.04.003. URL https://www.sciencedirect.com/science/article/pii/S0968000405000939

[49] J. Marles-Wright, R. J. Lewis, Stress responses of bacteria, Current Opinion in Structural Biology 17 (6) (2007) 755–760. doi:10.1016/j.sbi.2007.08.004. URL https://www.sciencedirect.com/science/article/pii/S0959440X0700111X

[50] T. M. Gruber, C. A. Gross, Multiple Sigma Subunits and the Partitioning of Bacterial Transcription Space, Annual Review of Microbiology 57 (Volume 57, 2003) (2003) 441–466, publisher: Annual Reviews. doi:10.1146/annurev.micro.57.030502.090913. URL https://www.annualreviews.org/content/journals/10.1146/annurev.micro.57.030502.090913

[51] S. Österberg, T. d. Peso-Santos, V. Shingler, Regulation of Alternative Sigma Factor Use, Annual Review of Microbiology 65 (Volume 65, 2011) (2011) 37–55, publisher: Annual Reviews. doi:10.1146/annurev.micro.112408.134219. URL https://www.annualreviews.org/content/journals/10.1146/annurev.micro.112408.134219

[52] R. A. Mooney, S. A. Darst, R. Landick, Sigma and RNA Polymerase: An On-Again, Off-Again Relationship?, Molecular Cell 20 (3) (2005) 335–345. doi:10.1016/j.molcel.2005.10.015. URL https://www.sciencedirect.com/science/article/pii/S1097276505016850

[53] K. L. Griffith, R. E. Wolf, Genetic Evidence for Pre-recruitment as the Mechanism of Transcription Activation by SoxS of Escherichia coli: The Dominance of DNA Binding Mutations of SoxS, Journal of Molecular Biology 344 (1) (2004) 1–10. doi:10.1016/j.jmb.2004.09.007. URL https://www.sciencedirect.com/science/article/pii/S0022283604011349

[54] W. Sung, M. S. Ackerman, M. M. Dillon, T. G. Platt, C. Fuqua, V. S. Cooper, M. Lynch, Evolution of the Insertion-Deletion Mutation Rate Across the Tree of Life, G3 Genes|Genomes|Genetics 6 (8) (2016) 2583–2591. doi:10.1534/g3.116.030890. URL https://doi.org/10.1534/g3.116.030890

[55] L. E. Williams, J. J. Wernegreen, Sequence Context of Indel Mutations and Their Effect on Protein Evolution in a Bacterial Endosymbiont, Genome Biology and Evolution 5 (3) (2013) 599–605. doi:10.1093/gbe/evt033. URL https://doi.org/10.1093/gbe/evt033

[56] J. W. Schroeder, W. G. Hirst, G. A. Szewczyk, L. A. Simmons, The Effect of Local Sequence Context on Mutational Bias of Genes Encoded on the Leading and Lagging Strands, Current Biology 26 (5) (2016) 692–697. doi:10.1016/j.cub.2016.01.016. URL https://www.sciencedirect.com/science/article/pii/S096098221600066X

[57] W. Sung, M. S. Ackerman, J.-F. Gout, S. F. Miller, E. Williams, P. L. Foster, M. Lynch, Asymmetric Context-Dependent Mutation Patterns Revealed through Mutation–Accumulation Experiments, Molecular Biology and Evolution 32 (7) (2015) 1672–1683. doi:10.1093/molbev/msv055. URL https://doi.org/10.1093/molbev/msv055

[58] A. Tanay, E. D. Siggia, Sequence context affects the rate of short insertions and deletions in flies and primates, Genome Biology 9 (2) (2008) R37. doi:10.1186/gb-2008-9-2-r37. URL https://doi.org/10.1186/gb-2008-9-2-r37

[59] A. P. W. Funnell, M. Crossley, Homo- and Heterodimerization in Transcriptional Regulation, in: J. M. Matthews (Ed.), Protein Dimerization and Oligomerization in Biology, Springer, New York, NY, 2012, pp. 105–121. doi:10.1007/978-1-4614-3229-67. URL

