## Supplementary Figure 1 for "Information Theory of Composite Sequence Motifs: Mutational and Biophysical Determinants of Complex Molecular Recognition"

---

---

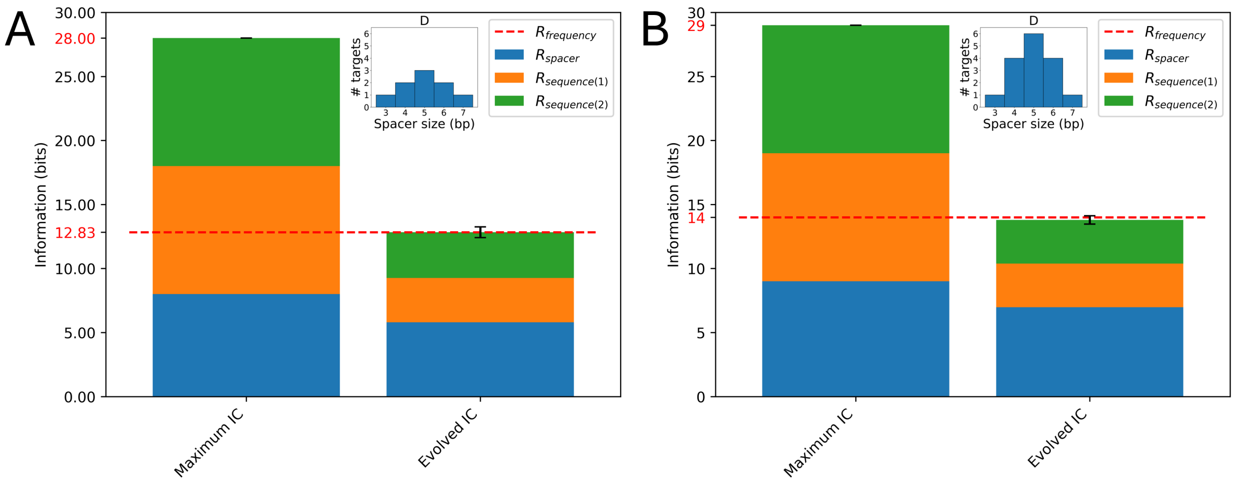


**Figure S1:** Distribution of evolved information for dyad-based systems ($n = 2$) in simulations using Gaussian connectors. The spacer size distribution ($D$) is shown as a histogram. Population size: 64. (A) Parameters: $\gamma = 9$, $G = 256$. (B) Parameters: $\gamma = 16$, $G = 512$.
